# AAV-mediated Expression of a Novel Conformational Anti-Aggregated α-Synuclein Antibody Prolongs Survival in a Genetic Model of α-Synucleinopathies

**DOI:** 10.1101/2022.11.30.518485

**Authors:** Matthias Düchs, Dragica Blazevic, Philipp Rechtsteiner, Cynthia Kenny, Thorsten Lamla, Sarah Low, Jimmy Savistchenko, Manuela Neumann, Ronald Melki, Tanja Schönberger, Birgit Stierstorfer, David Wyatt, Frederik Igney, Thomas Ciossek

**Affiliations:** Boehringer Ingelheim Pharma GmbH & Co KG, Biberach an der Riss, Germany; Boehringer Ingelheim USA, Ridgefield, Connecticut, USA; Institut Francois Jacob (MIRCen), CEA, CNRS, Fontenay-aux-Roses, France; Molecular Neuropathology of Neurodegenerative Diseases, German Center for Neurodegenerative Diseases and Department of Neuropathology, University Hospital of Tübingen, both Tübingen, Germany

## Abstract

Prion-like transmission of pathology in α-synucleinopathies like Parkinson’s disease or multiple system atrophy is increasingly recognized as one potential mechanism to address disease progression. Active and passive immunotherapies targeting insoluble, aggregated α-synuclein are already being actively explored in the clinic with mixed outcomes so far. Here, we report the identification of 306C7B3, a highly selective, aggregate-specific α-synuclein antibody with picomolar affinity devoid of binding to the monomeric, physiologic protein. 306C7B3 binding is Ser129-phosphorylation independent and shows high affinity to several different aggregated α-synuclein polymorphs, increasing the likelihood that it can also bind to the pathological seeds assumed to drive disease progression in patients. In support of this, highly selective binding to pathological aggregates in postmortem brains of MSA patients was demonstrated, with no staining in samples from other human neurodegenerative diseases.

To achieve CNS exposure of 306C7B3, an Adeno-Associated Virus (AAV) based approach driving expression of the secreted antibody within the brain of (Thy-1)-[A30P]-hα-Synuclein mice was used. Widespread central transduction after intrastriatal inoculation was ensured by using the AAV2HBKO serotype, with transduction being spread to areas far away from the inoculation site. Treatment of (Thy-1)-[A30P]-hα-Synuclein mice at the age of 12 months demonstrated significantly increased survival, with 306C7B3 concentration reaching 3.9 nM in the cerebrospinal fluid.

These results suggest that AAV-mediated expression of 306C7B3 has great potential as a disease-modifying therapy for α-synucleinopathies as it ensures CNS exposure of the antibody, thereby mitigating the selective permeability of the blood-brain barrier.

## Introduction

α-synucleinopathies, like Parkinson’s disease (PD), dementia with Lewy bodies (DLB) and multiple system atrophy (MSA), are relentless neurodegenerative diseases with no disease-modifying therapies for patients available to date. These diseases are characterized by intracellular deposits rich in an aggregated form of the 140 amino acid residue-long, intrinsically disordered protein α-synuclein (Shahmoradian et al., 2019; Oliveira et al. 2021). These deposits are known as Lewy bodies, Lewy neurites or oligodendroglial cytoplasmic inclusions and are regarded as histological hallmarks allowing definite classification of these diseases (Spillantini et al., 1997; Goedert et al., 2013), although exceptions exist for some rare familiar cases (reviewed by Schneider and Alcalay, 2017). α-synuclein had first been linked to PD based on the identification of a pathogenic mutation (Polymeropoulos et al., 1997) and since been confirmed to be genetically linked to PD based on additional point mutations as well as gene duplications and triplications causing highly penetrant autosomal forms of PD (Singleton et al., 2003; Chartier-Harlin et al., 2004). These familial point mutations as well as increased gene dosage of wildtype α-synuclein usually cause an early onset of disease combined with rapid disease progression. Overall incidence of mutation carriers is low, but genome wide association studies (GWAS) demonstrated an association between α-synuclein and the risk of developing disease even for idiopathic PD cases (Simon-Sanchez et al., 2009), with the association towards the SNCA gene coding for α-synuclein representing the strongest signal in PD GWAS studies identified so far (Brockmann et al., 2013).

The physiological function of α-synuclein is still poorly understood. The protein is highly enriched in presynaptic nerve terminals (Iwai et al., 1995). It has been suggested to play a role in presynaptic function (Vargas et al., 2014), and has been shown to allow SNARE complex assembly, a key component for membrane fusion between vesicles and presynaptic terminals during neurotransmission (Huang et al., 2019). However, genetic models with loss of α-synuclein expression present with only minor deficits (Abeliovich et al., 2000; Greten-Harrison et al., 2010; Burre et al., 2010). α-Synuclein aggregation leads to the loss of function of its physiological role (Sulzer and Edwards, 2019), but also to a gain of pathological function at the origin of PD, DLB or MSA. Recent work in experimental systems support the spread of misfolded, aggregated α-synuclein between neuronal cells (Luk et al., 2012a; Sacino et al., 2014; Peelarts et al., 2015; Sorrentino et al., 2017), potentially explaining the observed temporal and spatial progression of disease in affected patients (Braak et al., 2003), and the presence of Lewy bodies in juvenile grafts within the brain of PD patiens (Li et al., 2008; Kordower et al., 2008). These models assume that fibrillar α-synuclein is either secreted or passively released from neurons carrying intracellular aggregations, which spreads within the brain and – upon uptake – seeds the aggregation of endogenous α-synuclein in recipient neurons in a prion-like manner (reviewed by Brundin and Melki, 2017). An important element of this model is the fact that such seeds can be detected within cerebrospinal fluid (CSF) samples from patients (Manne et al., 2019; Fairfoul et al., 2016; Shahnawaz et al., 2017).

A potential prion-like cell-to-cell transmission of pathological aggregates of α-synuclein enabled the therapeutic concept of antibody-mediated blockage of disease progression in PD, DLB and MSA. This concept has been tested preclinically as well as clinically (summarized recently by Brundin et al., 2017), with mixed results, especially regarding beneficial effects in patients. Challenges to be considered in the clinical application of this concept include the problem of sufficient delivery of antibodies to the human brain (serum to plasma ratio of about 1:440; Reiber, 2001), especially since the blood-brain barrier seems to be only mildly perturbed in PD or MSA patients (Jesse et al., 2011; Song et al., 2011; see also actual exposure data in Jankovic et al., 2018 and Brys et al., 2019). In addition, some of the advanced immunotherapies (PDE03A, Poewe et al., 2021; PRX002, Jankovic et al., 2018) utilize antibodies with significant binding affinity towards non-aggregated, monomeric α-synuclein, a property that might be considered undesirable considering the high plasma exposure measurable in man (Barbour et al., 2008; Koehler et al., 2015). This potentially blocks the antigen binding site of these antibodies upon intravenous administration into patients as exemplified by the strong reduction in plasma α-synuclein upon PRX002 treatment (Schenk et al., 2017).

Herein, we describe the identification and characterization of a novel anti-α-synuclein antibody directed against the aggregated form of the protein that was counter-selected against monomeric α-synuclein. We detected sensitive and specific binding of our antibody to α-synuclein aggregates in post-mortem human brain tissue. Furthermore, the therapeutic efficacy of 306C7B3 was analyzed in a murine disease model with age-dependent development of pathology, based upon Adeno-Associated Virus (AAV) mediated antibody expression ensuring high central antibody exposure.

## Materials and Methods

### Expression and purification of recombinant monomeric human α-synuclein, and generation of aggregated polymorphs

Expression and purification of monomeric human and mouse α-synuclein used for immunizations and affinity determinations was performed as described by Danzer et al. (2017). To generate aggregated polymorphs, human wild type α-synuclein was purified as described by Ghee et al. (2005). Monomeric α-synuclein was aggregated into the fibrillar polymorphs “Fibrils” and “Ribbons” as described previously (Bousset et al., 2013) and into the polymorphs “Fibrils 91” and “Fibrils 65” as described in Makky et al. (2016). „Fibrils 110” that are made of human α-synuclein lacking 30 C-terminal amino-acid residues was generated as described (Shrivastava et al., 2020). The oligomeric forms OGA, ODA and O550 of α-synuclein were generated and purified as described (Pieri et al., 2016). All assemblies were assessed by Transmission Electron Microscopy (TEM) after adsorption onto carbon-coated 200 mesh grids and negative staining with 1% uranyl acetate using a Jeol 1400 transmission electron microscope and the images were recorded with a Gatan Orius CCD camera (Gatan, Pleasanton).

### Preparation of fibrillar α-synuclein phosphorylated at Ser129

For generation of pSer129 positive fibrillar α-synuclein preparations, monomeric human α-synuclein (2.5 mg/ml) was *in vitro* phosphorylated overnight at 30°C by incubation with purified PLK3 kinase (4.2 μg/ml, #PV3812, ThermoFisher) in a buffer consisting of 20 mM Hepes pH 7.4, 15 mM MgCl2, 1 mM Mg-ATP, and 2 mM DTT (protocol kindly provided by A. Oueslati and H. Lashuel). Successful phosphorylation was confirmed by mass spectroscopy indicating full pSer129 phosphorylation. Buffer exchange to enable fibril formation was performed using PD 10 spin columns. pSer129-phosphorylated fibrils were fragmented in a Covaris Focus ultrasonic device.

### Hybridoma screening

Hybridoma screening had been performed in collaboration with nanoTools Antikörpertechnik GmbH (Teningen, Germany). For immunization, fibrils of human α-synuclein were further purified to reduce contamination with residual monomeric protein. In brief, preparations were centrifuged at 100.000 g for 1h to sediment aggregated α-synuclein and the resulting pellet was resuspended three times with PBS followed by additional ultracentrifugations under identical conditions. Final preparations were fragmented in a classical ultrasonic water bath for 8h, aliquoted and stored at -80°C. After confirmation of bioactivity (see functional in vitro assay), mice were immunized with fibrils of human α-synuclein and underwent two additional immunizations 5 and 9 weeks after initial priming to boost antibody generation. Obtained hybridoma supernatants were tested in a Luminex bead-based ELISA for binding towards human fibrillar α-synuclein. Positive clones were counter-selected based on binding to monomeric human α-synuclein and confirmed to bind to murine fibrillar α-synuclein.

### Purification of antibodies

Recombinant antibodies (scIgG constructs) were purified after transient transfection of HEK-293 cells with the relevant expression constructs as described in the supplementary materials. Cell culture supernatants of transfected HEK-293 cells cultured in serum-free medium were collected 48h post-transfection and loaded onto a protein-A based column (AKTApurifier system, Mab Select SuRe 5ml pre-packed column). After extensive washing with PBS, 1 M NaCl, antibodies were eluted with 30 mM Sodium Acetate, pH 3.6 follow by size exclusion chromatography with 20 mM Na Citrate, 115 mM NaCl, pH 6.0 as running buffer. A similar protocol was used for the purification of native IgG from cultivated hybridoma clones.

### In vitro functional assay analyzing intracellular aggregation of α-synuclein

306C7B3 or other antibody preparations were pre-incubated at the indicated amounts (300 ng to 3 μg) with 50 μg monomeric human α-synuclein pre-filtered through Amicon Ultra-4 centrifugation filter tubes (100K, 4ml, #UFC 810024) for 3 h at room temperature to fully saturate monomer-binding antibodies. To each preparation 60 ng of fibrillar α-synuclein was added and further incubated for 30 minutes at room temperature to allow antibody binding to fibrillar α-synuclein. 1.5 million SH-SY5Y cells stably overexpressing human A53T mutated α-synuclein (Danzer et al., 2007) were resuspended in electroporation buffer R (NEON electroporation system, ThermoFisher) and added to the antibody-fibril preparation (final volume 100 μl). Cells were pulsed once at a voltage of 1200 V for 30 milliseconds. The electroporated cells were immediately diluted into 8 ml differentiation media for SH-SY5Y cells (Encinas et al., 2000) and seeded in collagen-IV coated flat-bottom 384 well microtiter plates amenable for high content analysis (Greiner) at a concentration of 10.000 cells per well in 80 μl total volume. Cells were incubated at 37°C for 2 days in a standard cell culture incubator. Percentage of cells carrying intracellular α-synuclein aggregates was analyzed after fixation in an InCell High Content Analyzer via immunofluorescence staining against pSer129 phosphorylated α-synuclein (anti-pSer129 antibody, Epitomics, #2014-1).

### Filter trap assay against different aggregate assemblies of human α-synuclein

Monomeric human α-synuclein and different fibrillar polymorphs (fibrils, ribbons, fibrils 65, fibrils 91, fibrils 110) and oligomeric species (on fibrillar assembly pathway α-synuclein oligomers O550, dopamine-stabilized ODA and glutaraldehyde-stabilized OGA oligomers) in the range of 20 pg to 200 ng were spotted on nitrocellulose filters (Ref Protran 0.45μm NC) using a slot blot filtration apparatus (GE Healthcare). The filters were blocked with skimmed milk and incubated with 306C7B3 IgG at 0.1 μg/ml. After extensive washing, binding of 306C7B3 was revealed using a goat anti-mouse secondary antibody with Super Signal ECL (Pierce #34096). The blots were imaged on a BioRad imager (Chemidoc MP imaging system/BioRad imagelab software).

### Affinity measurements

Antibody affinities were measured by surface plasmon resonance on a Biacore T2000 instrument. For monomeric α-synuclein, anti-murine IgG was immobilized on a CM5 chip. Test antibodies were captured using a 10 μg/ml antibody solution. Binding of monomeric α-synuclein was then analyzed in single-cycle kinetics with 5 concentrations ranging from 1.2 to 100 nM. Fibrillar α-synuclein prepared as described above was further purified by size exclusion chromatography (Superdex 75 10/300 GL column), and purity was confirmed by analytical HPLC (>95 %). Fibrils were immobilized on CM5 chips by amine coupling at 20 μg/ml in 10 mM Na-acetate buffer, pH 4.5. Binding of test antibodies was determined in single-cycle kinetics with 5 antibody concentrations ranging from 0.12 to 10 μg/ml. KD values were calculated using a 1:1 binding model adjusted for drift.

### Epitope determination

The identification of the epitope of selected antibodies was performed at PEPperPRINT GmbH (Heidelberg, Germany) utilizing the PEPperMAP epitope mapping technology. In brief, the full sequence of human α-synuclein was elongated at the C- and N-terminus with neutral GSGSGSG linkers and translated into an array of 15 amino acid peptides with a peptide overlap of 14 amino acids. The resulting peptide microarray containing 140 different peptides was printed in duplicate and framed by Flag (DYKDDDDKGG) and HA (YPYDVPDYAG) control peptides. Peptide microarrays were incubated with the antibody in question, followed by washing steps and staining with a secondary goat anti-mouse antibody conjugated with fluorophore dyes. After analysis for signal intensity, an intensity map based on average median foreground intensities was generated and peptides contributing to antibody binding calculated.

### Human postmortem tissue

Sections from paraffin-embedded formalin-fixed human brain tissue samples were obtained from the brain banks affiliated with the University of Tübingen and the DZNE. Consent for autopsy was obtained from probands or their legal representative in accordance with local institutional review boards. The cohort of α-synucleinopathy cases consisted of six cases with Lewy body diseases (PD, DLB) and three cases with MSA. Control cases for immunohistochemistry included frontotemporal lobar degeneration with TDP-43 pathology (FTLD-TDP; n=4), FTLD with FET pathology (FTLD-FET; n=1), sporadic amyotrophic lateral sclerosis with TDP-43 pathology (ALS; n=2), progressive supranuclear palsy (PSP; n=2), Alzheimer’s disease (AD; n=2), and neurologically healthy controls (n=2).

### Immunohistochemistry

Immunohistochemistry on human postmortem tissues was performed on 3 μm thick sections of formalin fixed, paraffin-embedded (FFPE) tissues from neuroanatomical regions with robust pathology in the respective diseases using the Ventana BenchMark XT automated staining system with the iVIEW DAB detection kit (Ventana). 306C7B3 was used at 1:10000 dilution with heat pretreatment (boiling for 32 min in CC1 buffer, Ventana). Other antibodies included anti-pS409/410-TDP-43, clone 1D3 (1:500, Neumann et al., 2009), anti-ptau, clone AT8 (ThermoFisher, 1:500), anti-α-synuclein, clone 4D6 (Origene, 1:1000), anti-beta-Amyloid, clone 4G8 (Covance, 1:6000) and anti-FUS (Bethyl Laboratories, 1:400).

In addition, manual staining was done on selected human case and mouse tissue. The tissue sections were deparaffinized in a standard series of xylol and alcohol. Pretreatments varied depending on the individual antibodies employed. For GFP (rabbit polyclonal anti-GFP, Abcam, ab290), sections were incubated for 10 minutes in enzyme buffer (Bond Enzyme Pretreatment Kit, Leica Biosystems) at 37°C to assist epitope retrieval. 306C7B3 stainings were done with enzyme pretreatment (Bond Enzyme Pretreatment Kit, Leica Biosystems) at 37°C for 5 minutes. Syn-1 stainings (clone 42, BD Biosciences) were done at a dilution of 1:1000. For pSer129 α-synuclein staining (EP1536Y rabbit anti-α-synuclein pSer129 antibody, Epitomics, diluted 1:1000), no pretreatment was required. Color development was performed with classical DAB (3,3’-diaminobenzidine) staining (Bond Polymer Refine Detection Kit, Leica Biosystems.

### AAV Production and Quantification

Expression plasmids for AAV production were generated by gene synthesis (GeneArt, ThermoFisher) and subcloned into the pAAV plasmid (AAV Helper-Free System, Agilent). A plasmid map for the scIgG-306C7B3 expression construct is shown in the Supplementary Materials. Viral stocks were generated as described by Strobel et al. (2021) using polyethylenglycol precipitation and iodixanol gradient centrifugation of HEK293 cell lysates. AAVs were dissolved in AAV formulation buffer (PBS, 1 mM MgCl2, 2.5 mM KCl, 10% glycerol, 0.001% Pluronic F-68, pH 7.4) and stored at -80°C after being sterile-filtered. Titer determination with quantitative PCR was done on extracted viral DNA according to Strobel et al. (2021) with PCR primers directed against the promoter region of the expression constructs.

### In vivo experiments in (Thy-1)-h[A30P]-α-synuclein mice

All animal experiments were conducted in an AAALAC-accredited facility (Association for Assessment and Accreditation of Laboratory Animal Care International) in accordance with the EU Directive 2010/63/EU for animal experiments. Authorization was granted by the Animal Care Commission of the government of Baden Wuerttemberg, Germany (Regierungspräsidium Tübingen). Mice were given food and water ad libitum, while room temperature and humidity were maintained at 22°C ± 2°C and 55% ± 10%, respectively.

For the described experiments homozygous (Thy-1)-h[A30P]-α-synuclein mice (Kahle et al., 2000; Neumann et al., 2002; kindly donated by Philipp Kahle) extensively backcrossed (>10 generations) into C57Bl6/J background were used. Phenotype development in this transgenic model is dose dependent with earlier onset of pathology in homozygous compared to heterozygous animals. To ensure homozygosity of the transgene, genotyping was performed on tail biopsies taken from animals after weaning. Genomic tail DNA was purified using QiaAmp DNA Mini Kits and a Qiacube robot (Qiagen), according to the instructions of the manufacturer. Briefly, tail snips were digested overnight at 56°C with shaking in tissue-lysis buffer containing proteinase K, followed by purification with the provided spin columns. Initially, genotyping was performed with quantitative PCR able to distinguish hemizygous from homozygous animals as described by Scudamore et al. (2018). Later genotyping was done with classical PCR based on primers spanning the integration site of the transgene, resulting in amplified fragments of 287 bp for the transgene allele and 340 bp for the wildtype allele (Gentzel et al., 2021).

### Stereotactic injection into the striatum of (Thy-1)-h[A30P]-α-synuclein mice

AAVHBKO vectors expressing either scIgG-306C7B3 or a control antibody without a known target in mice (anti-FITC scIgG) were stereotactically injected (1 μl per injection, total titers as described in the figures) bilaterally into the striatum of (Thy-1)-h[A30P]-α-synuclein male and female mice at an age of 12 months (n=16 per group). All surgical procedures were performed using aseptic techniques. Briefly, medetomidin, midazolam, fentanyl (0,5/5/0,05 mg per kg intraperitoneally) was used for anesthesia. After onset of anesthesia, the head was shaved, area disinfected, local anesthetic lidocaine applied, and the animal transferred to a stereotactic head frame. Body temperature was controlled and animals were kept on a heat pad. A midline anterior posterior incision of the scalp was used to access the skull and lambda and bregma were used for anterior-posterior (+0,1/+0,1 cm), dorsal-ventral (−0,37/-0,36 cm) and medial-lateral (+0,2/-0,2 cm) orientation for bilateral application. Burr holes were drilled in the appropriate locations and 1 μl of fluid with 200 nl/min was injected into the striatum using a microliter syringe (Hamilton, Reno, Nevada). Injection needle was kept in place for 5 min before careful retraction from the injection side. After cleaning of the skull and stitching of the scalp, Atipamizol, Flumazenil, Naloxon (2,5/0,5/1,2 mg per kg subcutaneous) were given to antagonize anesthesia. For post operative care meloxicam (1 mg per kg subcutaneous) was used.

Animals were constantly monitored for disease progression as indicated by the loss of righting reflex (Scudamore and Ciossek, 2018), triggering euthanasia of the affected animals. Brain tissue as well as a final blood sample for exposure measurements were prepared and further analyzed as described.

### Vector genome quantification

Vector genomes present in each treated animal was quantified after preparation of genomic DNA from one brain hemisphere with the AllPrep96 DNA/RNA Kit (Qiagen). qPCR with primers and a probe directed against the hSyn promoter sequence (forward 5’-ATGAGTGCAAGTGGGTTTTAGGA-3’, reverse 5’-CCCTCCCCCTCTCTGATAGG-3’, probe 5’-CGACCCCGACCCACTGGACAAG-3’) was performed to confirm efficient and dose-dependent transduction.

### CSF sampling

CSF was collected from the cisterna magna. Therefore, animals were fixated and anesthetized as described for stereotactic injections. Briefly, after application of local anesthetics the head of the animal was bend and the skin at the neck of animal opened with an incision on the base of the skull. Tissue was bluntly removed to access cisterna magna via occipital membrane. A microliter syringe (Hamilton, Reno, Nevada) was guided via the stereotaxic frame and a micropump (Hamilton, Reno, Nevada) was used to withdraw fluid at a speed of 1000nl/min.

### ELISAs to measure 306C7B3 and anti-FITC antibodies

An MSD-based ELISA was established to measure exposure of 306C7B3 in plasma and CSF mouse samples, similar to the description provided in Strobel et al. (2020). Briefly, MSD standard plates (MSD #L15XA-1) were coated overnight with 100 ng fibrillar human α-synuclein (diluted to 30 μl per well with PBS) at 4°C overnight. After washing to remove unbound fibrillar α-synuclein, plasma or CSF probes were incubated in 25 μl for 1 hour at room temperature (diluted with PBS containing 5% Amersham ECL Blocking Agent, GE Healthcare RNP2125), followed by additional washing steps and incubation with an anti-mouse secondary antibody conjugated with the MSD Sulfo-tag for detection. A similar protocol was established to measure the control anti-FITC antibody utilizing FITC-conjugated BSA (A23015, Molecular Probes) to capture the antibody (Strobel et al., 2020).

### Statistical analysis

The in vivo data was statistically analyzed based on a Cox regression model with time to death as dependent variable and treatment, gender and cause of death as independent factors. The statistical evaluation was prepared using the software SAS Version 9.4 (SAS Institute Inc., Cary, North Carolina, USA).

## Results

### Hybridoma screening for aggregate specific anti-human α-synuclein antibodies

Bacterially expressed and highly purified human α-synuclein was aggregated in vitro to pre-formed fibrillar α-synuclein (PFF), based upon established methods (Polinski et al, 2018). Care was taken to confirm biological activity of the obtained preparations by characterization in an SH-SY5Y based cellular assay, in which their potential to drive intracellular aggregation of human α-synuclein is measured by the appearance of mesh-like, pSer129-positive human α-synuclein aggregates. Active preparations underwent further biophysical purification to deplete monomeric α-synuclein as much as possible. Mice were repeatedly immunized with these aggregated preparations of human α-synuclein and isolated hybridomas screened for clones with high binding activity towards aggregated α-synuclein, combined with low to no binding to monomeric human α-synuclein. Selected clones were further characterized as depicted in supplementary figure 1.

### Characterization of 306B7C3

As shown in Figure 1A (left panel), the hybridoma clone 306C7B3 was finally selected based on its strong and almost irreversible binding to aggregated human α-synuclein with a calculated K_D_ of 15 pM and a k_off_ of less than 10^−5^/s, as determined in a surface plasmon resonance assay. Importantly, no binding to monomeric human α-synuclein could be measured (Figure 1A, right panel). 306C7B3 was further characterized for its functional activity regarding seeding of endogenous α-synuclein (aggregation indicated by Ser129 phosphorylation) in the above-described cellular assay (Figure 1C). Prior to electroporation of the SH-SY5Y cells overexpressing A53T-mutated α-synuclein, the antibody was pre-incubated with a large molar excess of monomeric human α-synuclein to fully block monomer-binding ability, followed by the addition of pre-formed fibrillar α-synuclein as seeds for intracellular aggregation (assay adapted from Bousset et al., 2013). 306C7B3, purified from its hybridoma supernatant, showed strong dose-dependent blockage of intracellular α-synuclein aggregation as analyzed by high content analysis of pSer129-positive intracellular aggregates (Figure 1D, “hybridoma”). The primary amino acid sequence of 306C7B3 was determined by sequencing the hybridoma clone and 306C7B3 converted to a single-chain antibody (scIgG) by joining the light and heavy antibody chains with a covalent linker as shown in supplementary Figure 2A. This allowed the expression of 306C7B3 from a single open reading frame (scIgG-306C7B3). This construct, purified from HEK293 cells transiently transfected with the corresponding expression construct (similar to the one shown in supplementary Figure 2C but carrying an ubiquitous CMV promoter instead of the depicted neuron-specific human synapsin promoter), is as active as the 306C7B3 purified from the hybridoma (Figure 1D, “scIgG”).

**Figure 1.**
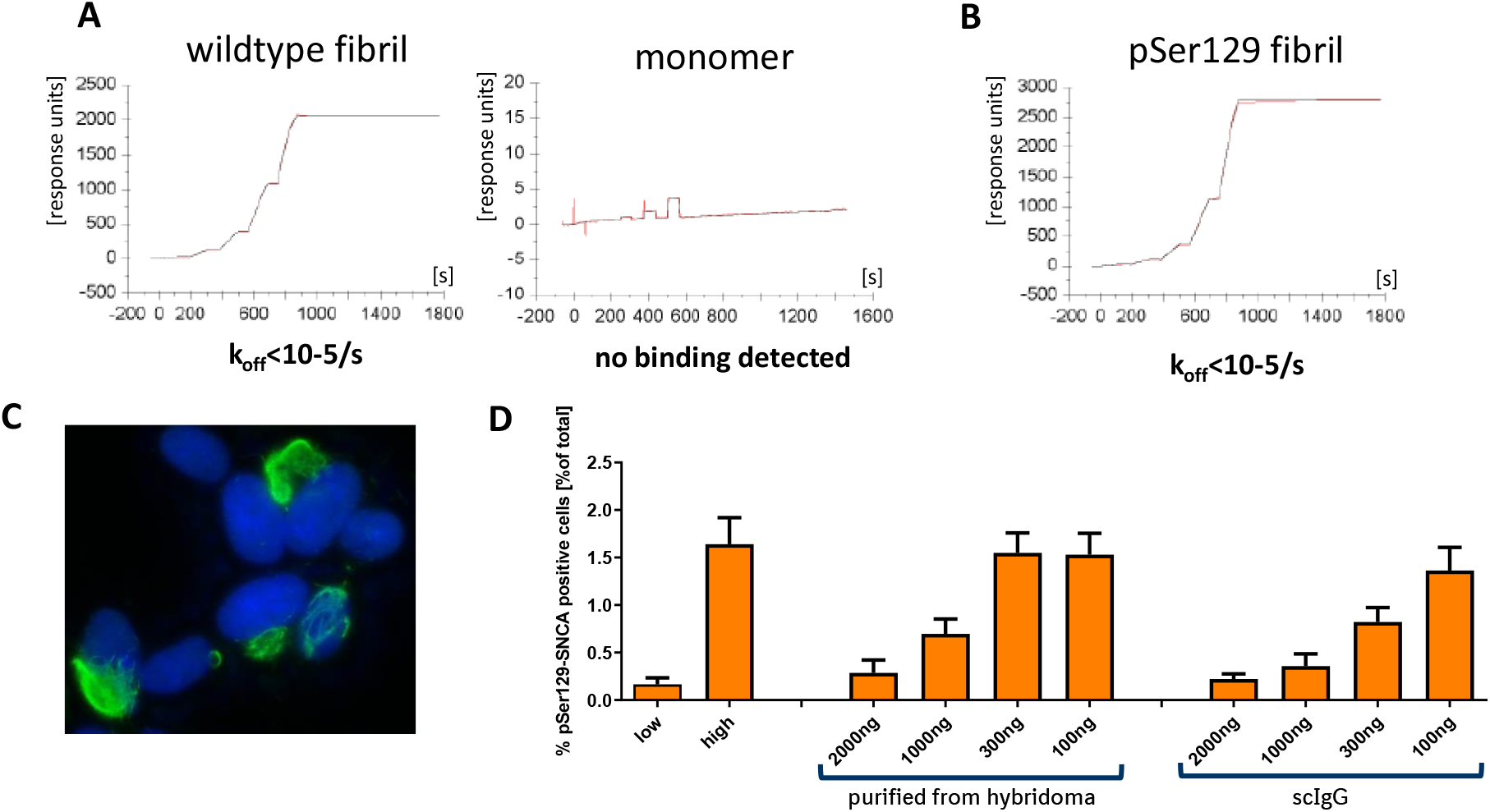
In vitro characterization of 306C7B3. ***A***, Surface plasmon based affinity determination of 306C7B3 for fibrillar human α-synuclein (left) as well as for monomeric human α-synuclein (right). ***B***, Similar measurement as in A, but using *in vitro* Ser129-phosphorylasted fibrillar human α-synuclein for the affinity determination. ***C***, Immunofluorescence staining of SH-SY5Y cells stably overexpressing A53T-mutated α-synuclein two days after electroporation with fibrillar α-synuclein. Intracellular mesh-like aggregates of pSer129 positive α-synuclein (green, Epitomics pSer129 antibody) are used for quantitative determination of functional activity of anti-α-synuclein antibodies (blue: DAPI staining). ***D***, Functional characterisation of 306C7B3. Increasing amounts of purified antibody derived either from the original hybridoma clone or from a scIgG-based expression construct were preincubated with an excess of monomeric human α-synuclein, followed by addition of PFF α-synuclein and electroporation into SH-SY5Y cells stably overexpressing human A53T α-synuclein. High content based analysis of cells stained for pSer129-positive intracellular aggregates demonstrates efficient inhibition of intracellular seeding by IgG-306C7B3 purified from the original hybridoma clone as well as single chain IgG 306C7B3 (scIgG) derived from an expression construct.

### Epitope mapping of 306C7B3

The epitope of 306C7B3 was determined via PEPperMAP epitope mapping as shown in Figure 2A. Control antibodies like Syn-1 (Figure 2B) showed strong and sharp signals at an antibody concentration of 1 μg/ml, with similar data obtained for e.g., 9E4 (Masliah et al, 2005, data not shown). However, no such staining could be observed with 306C7B3 at this antibody concentration. Only by raising the antibody concentration tenfold (Figure 2, red line), a broad area of 306C7B3 binding to C-terminal human α-synuclein 15-mer peptides could be observed, defining its epitope (“YQDYEPE”, amino acids 133 – 139; as calculated based upon the individual binding intensities to the 15mer α-synuclein peptides). This, together with the absence of binding to monomer (Figure 1A), suggests that 306C7B3 rather recognizes a conformational epitope human α-synuclein adopts within PFFs, and not to a linear amino acid epitope.

**Figure 2.**
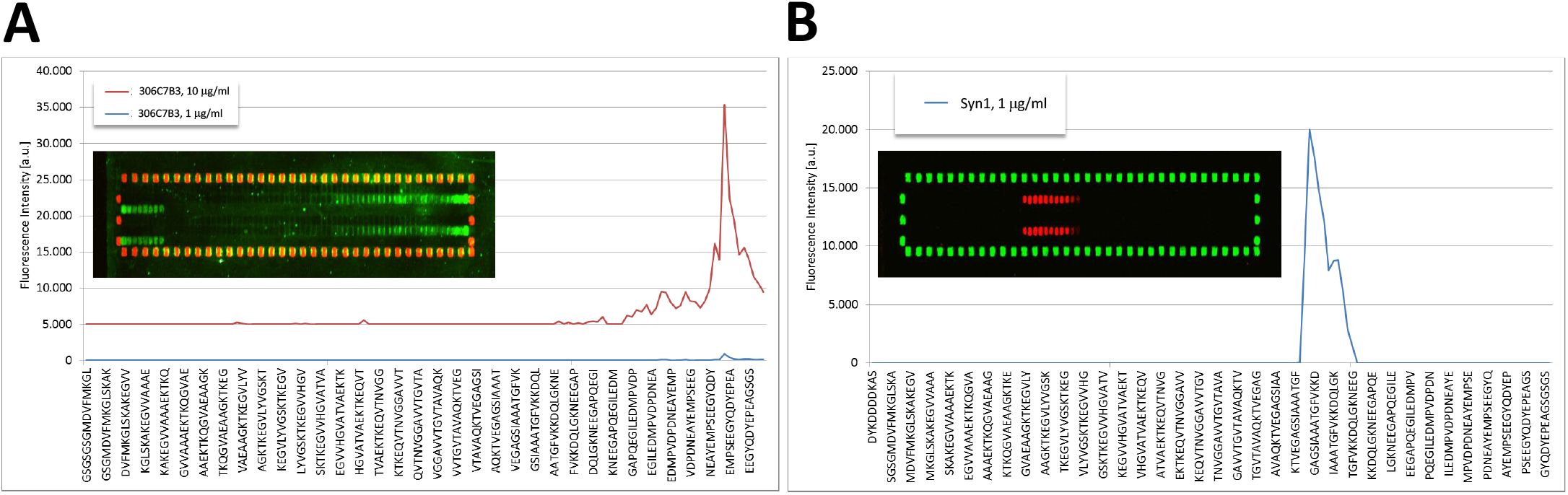
PEPperMAP Epitope mapping. ***A***, Either 1 μg/ml (blue line) or 10 μg/ml (red line) of IgG 306C7B3 were incubated with membranes spotted with 140 different 15mer peptides (each peptide printed in duplicate to be present in the top and bottom part of the array), representing the full human α-synuclein sequence with each peptide having a 14 amino acid overlap with the preceeding peptide with human α-synuclein sequence being elongated by neutral GSGSGSG linkers to avoid truncated peptides. Inset shows microarray stained with 10μg/ml 306C7B3 (green: 306C7B3 signal, red: HA signal). Outer spots correspond to control peptides used to orient the microarray. Some cross-reactivity against the control peptides is observed for 306C7B3 at this high antibody concentration. Figure shows signal intensity obtained for individual peptide spots, identifying the epitope of 306C7B3 as being „YQDYEPE” (amino acids 133-139 of full-length human α-synuclein) located in the C-terminus of α-synuclein with overall low binding intensity towards the microarray spotted α-synuclein peptides. ***B***, Similar approach to determine the epitope of Syn1 at a concentration of 1 μg/ml (blue line, red: Syn1 signal, green: HA signal), identifying the epitope of Syn1 as being „AATGFVKK” (amino acids 90-97 of full-length human α-synuclein). Note that the exemplified peptide sequences do not reflect all tested 15mer peptides for better visualisation – identical peptide arrays were used for the analysis of the different antibodies.

### 306C7B3 binding to structurally distinct human α-synuclein assemblies

306C7B3 was identified by using human α-synuclein PFFs aggregated in a buffer at physiological pH as used for many *in vitro* as well as *in vivo* studies (Luk et al., 2012a; Polinski et al., 2018). The extent to which 306C7B3 binds to structurally well characterized pure fibrillar and oligomeric α-synuclein polymorphs was assessed by filter trap measurements. Increasing amounts of the fibrillar polymorphs fibrils and ribbons (Bousset et al., 2013), fibrils 65 and fibrils 91 (Makky et al., 2016), fibrils 110 (Shrivastava et al., 2020), on fibrillar assembly pathway oligomers as well as dopamine or glutaraldehyde stabilized oligomers (Pieri et al., 2016) were spotted on nitrocellulose membrane and the binding of 306C7B3 quantified (Figure 3). 306C7B3 bound to all tested assemblies except fibrils 110, consistent with the absence of the α-synuclein C-terminal amino acid residues in fibrils 110. 306C7B3 barely bound to monomeric human α-synuclein. Finally, decreased binding was observed to dopamine or glutaraldehyde stabilized oligomers.

**Figure 3.**
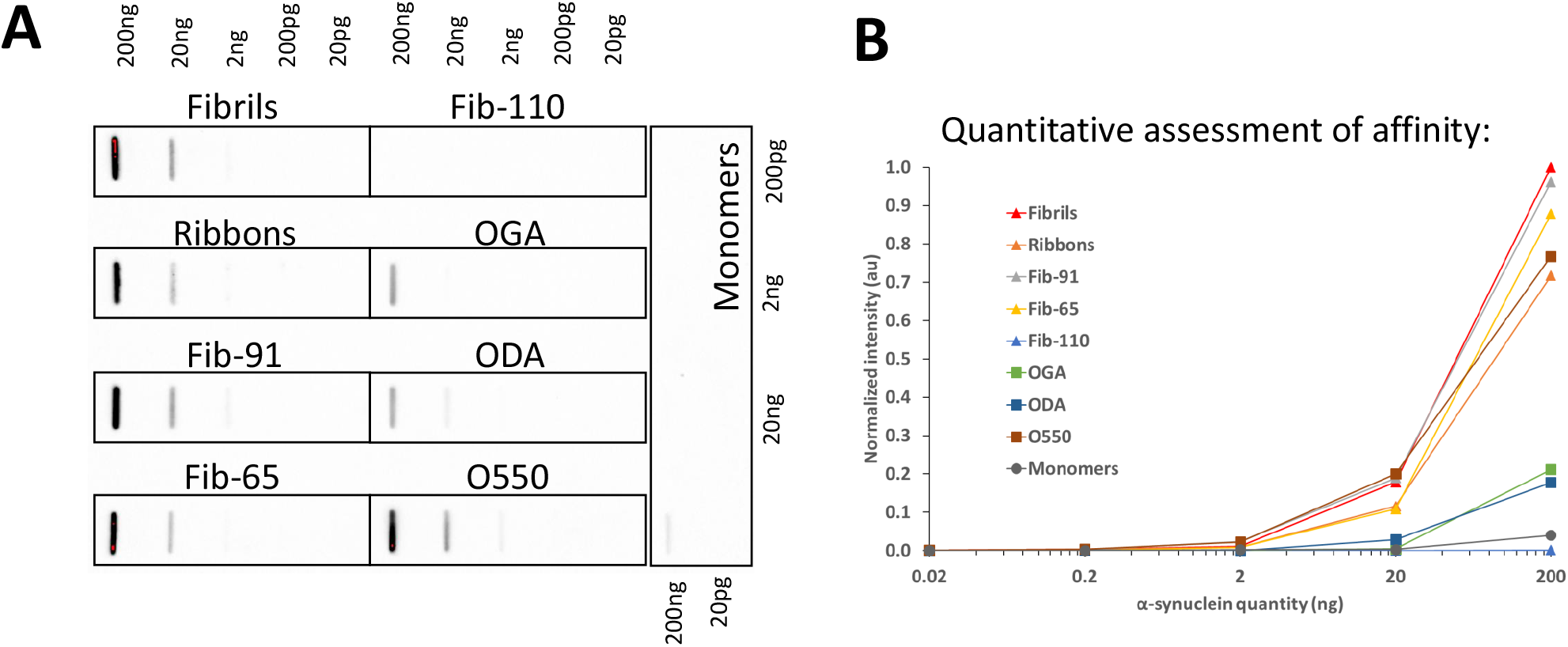
Filter trap assessment of 306C7B3 binding towards different *in vitro* generated assemblies of human α-synuclein. Fibrils, ribbons, fibrils 65, fibrils 91, fibrils 110, on fibrillar assembly pathway oligomers (O550), dopamine stabilized (ODA) and glutaraldehyde stabilized (OGA) oligomers were spotted in increasing amounts on nitrocellulose filters and tested with 306C7B3. ***A***, Images obtained after ECL based visualisation. ***B***, Quantification of signals using a Chemidoc MP imaging system.

One of the most prominent post-translational modifications affecting α-synuclein in patient brains and in *in vitro* and *in vivo* models is the phosphorylation of Ser129. This residue lies close to the epitope of 306C7B3 as determined in the PEPperMAP epitope assessment and may interfere with 306C7B3 binding. A surface plasmon resonance assay comparing binding of 306C7B3 to non-phosphorylated as well as Ser129-phosphorylated, aggregated human α-synuclein showed that Ser129 phosphorylation does not affect 306C7B3 binding (Figure 1B).

### Binding of 306C7B3 to pathological α-synuclein-rich deposits in human α-synucleinopathies

We next assessed the binding of 306C7B3 to pathological α-synuclein-rich deposits using postmortem tissue from patients suffering from MSA. As controls, the Syn-1 antibody known to display aggregation-independent α-synuclein binding, and an anti-pSer129-specific α-synuclein antibody were used. Syn-1, in addition to clearly identifying the pathological glial cytoplasmic inclusions in the white matter tracts of the cerebellum, also markedly stained monomeric α-synuclein in the neuropil of the molecular layer, corresponding to the presynaptic localization of monomeric α-synuclein (Figure 4A). This prominent staining for the physiological α-synuclein is absent in the immunohistochemical stainings with either 306C7B3 or the pSer129-specific antibody (Figure 4B and 4C), demonstrating a strong and aggregate-specific binding of 306C7B3 to human pathological α-synuclein-rich deposits. Strong immunoreactivity of cortical and subcortical neuronal and oligodendroglial inclusions could also be demonstrated in additional affected brain regions of MSA patients (supplementary Figure 3) as well as strong labeling of Lewy pathology in PD/DLB cases (data not shown). Furthermore, 306C7B3 binding was specific to α-synuclein deposits with no immunoreactivity observed to other amyloid deposits characteristic of neurodegenerative diseases such as TDP-43 inclusions in FTLD-TDP and ALS, FUS inclusions in FTLD-FET, tau pathology in PSP and AD, as well as Abeta in AD (Figure 5).

**Figure 4.**
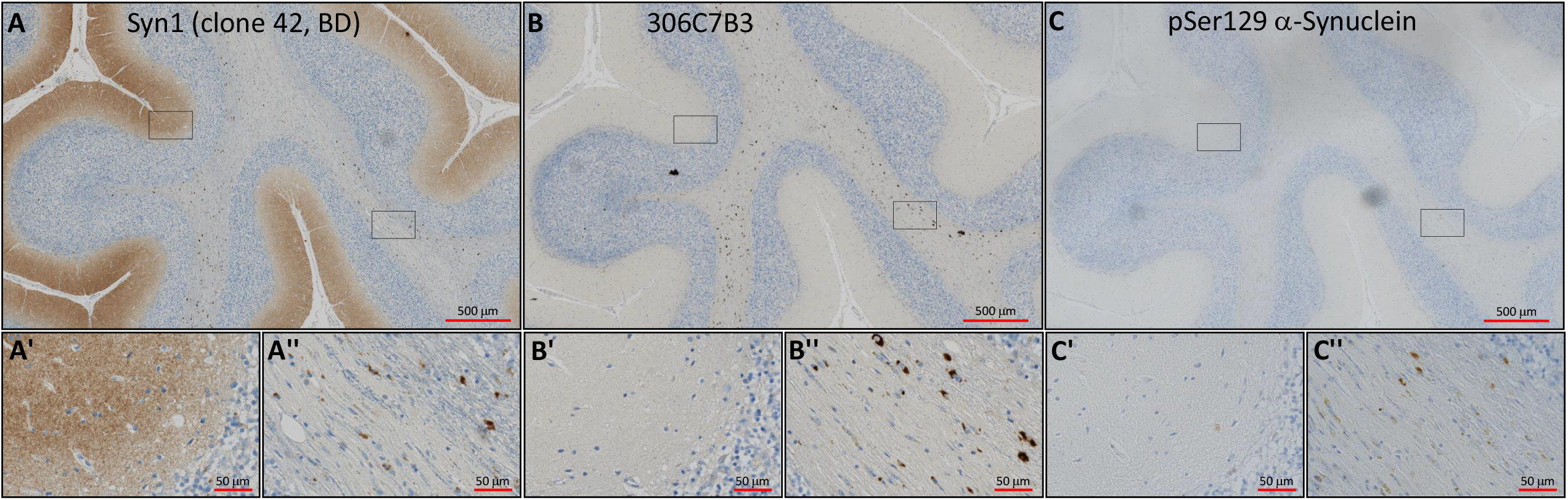
Immunohistochemical staining of adjacent sections of cerebellar tissue derived from a patient with Multiple System Atrophy (age at death 74 years, Braak&Braak stage I-II). ***A***, Staining performed with the Syn1 antibody detecting monomeric α-synuclein in the molecular layer as well as aggregated forms of α-synuclein in the white matter fiber tracts. ***B***, 306C7B3 staining for aggregated α-synuclein in the white matter fiber tracts only. ***C***, Control staining against pSer129-positive α-synuclein aggregates demonstrating a similar staining pattern compared to 306C7B3. ***A*’, *B*’, *C*’**, enlargement of the grey matter area indicated above (black box on the left, approximate location). ***A*’’, *B*’’, *C*’’**, enlargement of fiber tract staining in the white matter as indicated above (black box on the right, approximate location).

**Figure 5.**
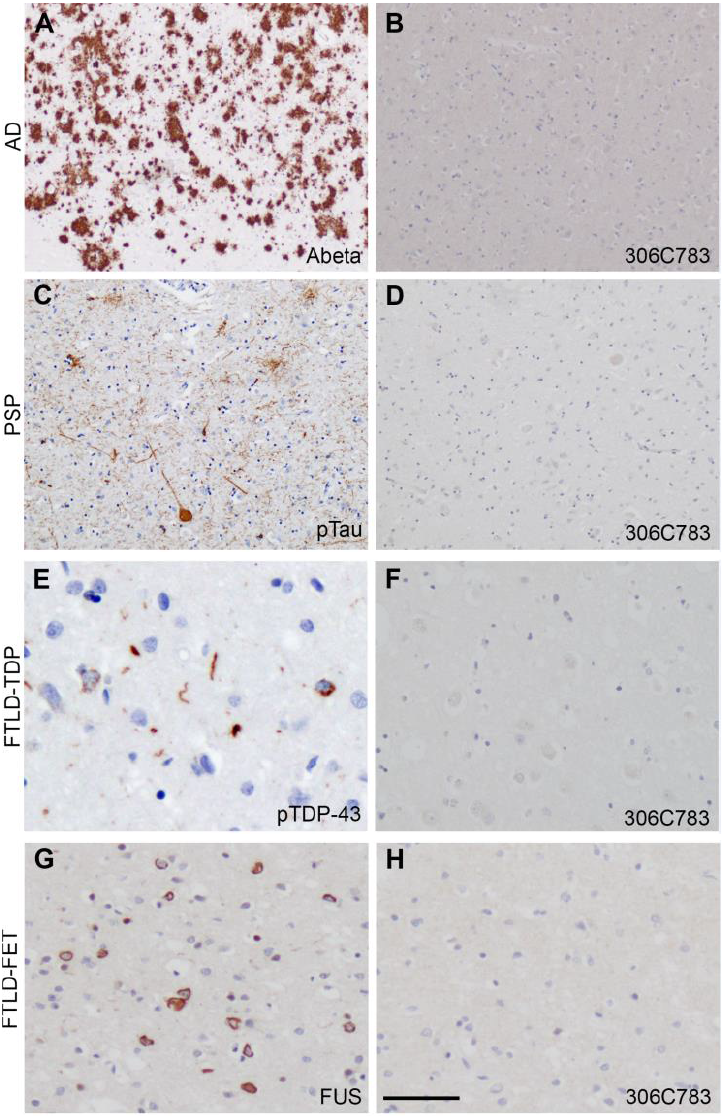
306C7B3 does not label other pathological aggregates. Immunohistochemical analysis of 306C7B3 revealed no labeling of pathological deposits characteristic for other neurodegenerative diseases, including Abeta-positive senile plaques in Alzheimer’s disease (AD, A and B), tau-positive tangles and tufted astrocytes in progressive supranuclear palsy (PSP, C and D), TDP-43-positive inclusions in frontotemporal lobar degeneration with TDP-43 pathology (FTLD-TDP, E and F), or FUS-positive inclusions in frontotemporal lobar degeneration with FET pathology (FTLD-FET, G and H). Immunohistochemistry on formalin fixed paraffin embedded adjacent tissue sections for 306C7B3 (B, D, F, H) and Abeta (A), pTau (C), pTDP-43 (E), or FUS (G). Scale bar 200μm (A-D), 40μm (E-H).

Based on the affinity and epitope data, the functional activity in the *in vitro* aggregation assay, the binding to different α-synuclein pathogenic assemblies, and the specific and sensitive staining of human pathological α-synuclein deposits in brain sections of MSA patients, we conclude that 306C7B3 is a highly aggregate specific anti-α-synuclein antibody devoid of binding towards monomeric α-synuclein, able to block the seeding activity of exogenous α-synuclein fibrils *in vitro*.

### AAV-mediated Antibody expression – selection of AAV serotype

Antibody delivery to the central nervous system is severely hampered by the blood brain barrier, restricting central exposure of full-length antibodies (as measured in the cerebrospinal fluid, CSF) to about 0.2% of the plasma exposure (Reiber, 2001). No major breakdown of the blood brain barrier has been observed in PD or MSA patients (Jesse et al., 2011; Song et al., 2011). To enable significant CSF exposure, we opted for Adeno-Associated Virus (AAV) mediated central transduction with an expression construct for 306C7B3, making use of the scIgG-based, single open-reading frame construct exemplified in the supplementary Figure 2A, an approach also termed “vectorized immunization” (Liu et al., 2016). To establish the route of administration, and to select a suitable AAV serotype, a pilot experiment comparing stereotactic delivery of AAV2 and AAV2HBKO, a variant of AAV2 devoid of heparan sulfate proteoglycan binding (Naidoo et al., 2018), into the murine striatum was performed (Figure 6). Both viral preparations carried an identical transgene coding for GFP under the control of the ubiquitous CMV promoter. Intrastriatal inoculation of the AAV2 based preparation at a total dose of 2.8×10^8^ vector genomes (vg) caused focal, high intensity staining with limited spread of transduction from the site of inoculation. In addition, the center of the transduced striatal area close to the inoculation site contained cellular infiltrates indicative of an inflammatory reaction (Inlets A’ and A’’ in Figure 6). In stark contrast, AAV2HBKO transduced striatum (Figure 6B) showed widespread transduction covering a large portion of the striatum extending into cortical areas in the vicinity of the striatum. No cellular infiltrates were observed in AAV2HBKO-inoculated striata. Therefore, the AAV2HBKO serotype was selected for further experiments, allowing for a larger area of transduction with a limited potential to drive inflammatory reactions.

**Figure 6.**
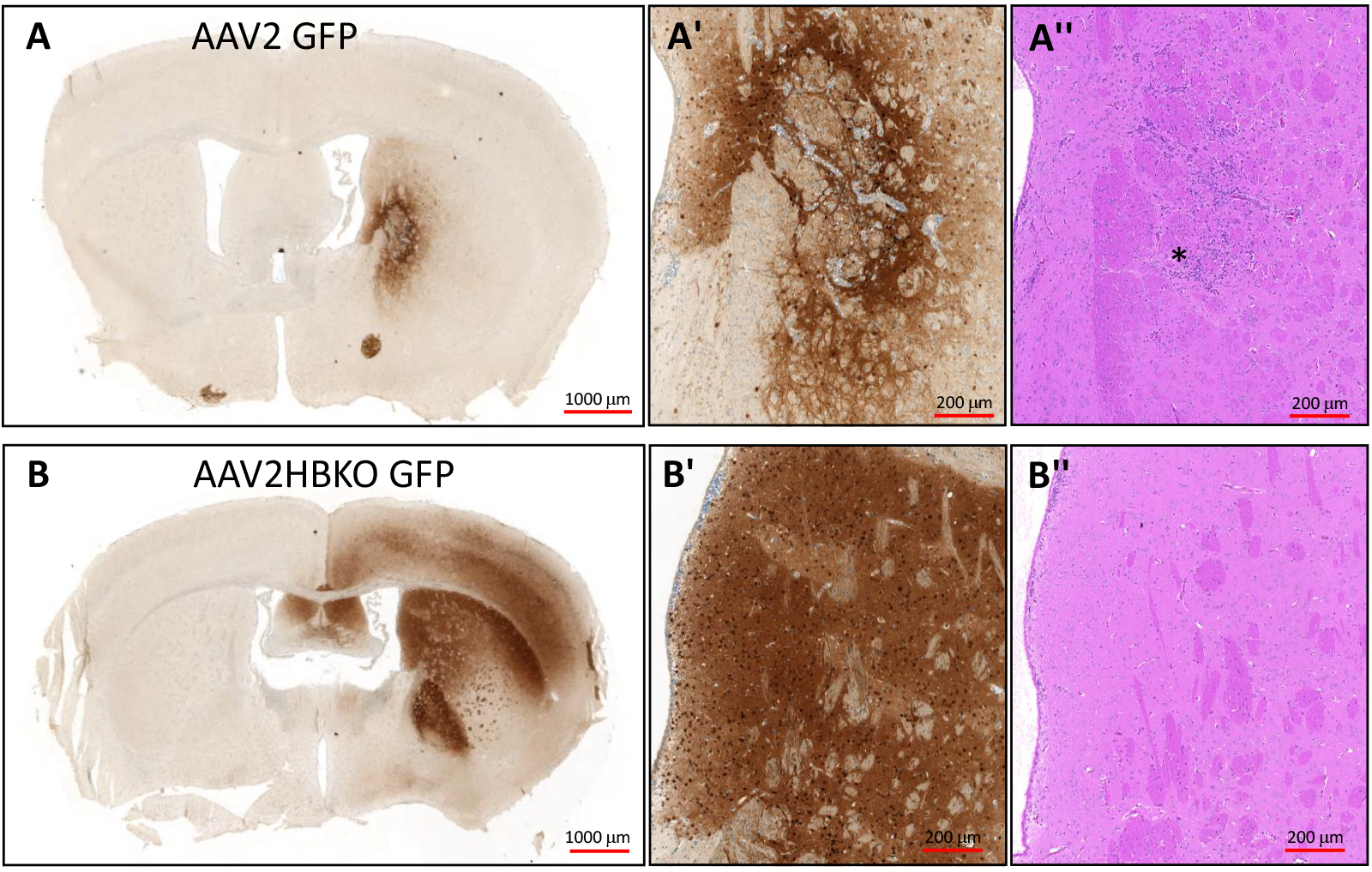
Expression of GFP in the murine striatum after AAV inoculation. One μl of self-complementating (sc) AAV preparations (total of 2.8×10^8^ vector genomes) coding for GFP under the control of the ubiquitous CMV-promoter were stereotactically inoculated into the murine striatum. Immunohistochemical analysis was done 21 days after virus administration. ***A***, AAV2 based transduction with limited spread of the viral preparation (anti-GFP IHC). ***B***, Significant spread of the viral transduction observed with AAV2HBKO based viral preparations (anti-GFP IHC). ***A*’, *B*’**, enlargement of the transduced striatum (anti-GFP IHC). ***A*’’, *B*’’**, H&E staining of adjacent sections in the area of the enlargements (striatum), demonstrating cellular infiltrates in the inoculation area of the AAV2-GFP viral preparation (marked by the asterix), no such infiltrations were observed in AAV2HBKO treated animals.

### In vivo α-synucleinopathy model – AAV-mediated expression of scIgG-306C7B3

(Thy-1)-[A30P]-hα-Synuclein mice, a transgenic model expressing the familial Parkinson’s disease A30P mutation of α-synuclein under the control of the neuronal Thy-1 promoter, develop α-synuclein pathology in a time-dependent manner (Neumann et al, 2002; Figure 7A). Previous experiments based on the quantification of pSer-129 α-synuclein load within brain lysates, or extent of central α-synuclein aggregation within the CNS (pSer129 α-synuclein positive staining by immunohistochemistry), exhibited significant variations precluding quantitative analysis (data not shown). We therefore designed a survival study focused on the analysis of the loss of righting reflex as an indicator of severe central α-synucleinopathy (Scudamore and Ciossek, 2018) to circumvent the variability in onset and aggregate load in this model, to characterize potential effects of 306C7B3 *in vivo*. Figure 7B shows survival of non-treated control animals. Animals euthanized due to the loss of righting reflex were confirmed to be homozygous for the α-synuclein transgene (supplementary Figure 4) as well as to carry strong central α-synuclein pathology as analyzed by pSer129-positive α-synuclein immunohistochemistry (supplementary Figure 5).

**Figure 7.**
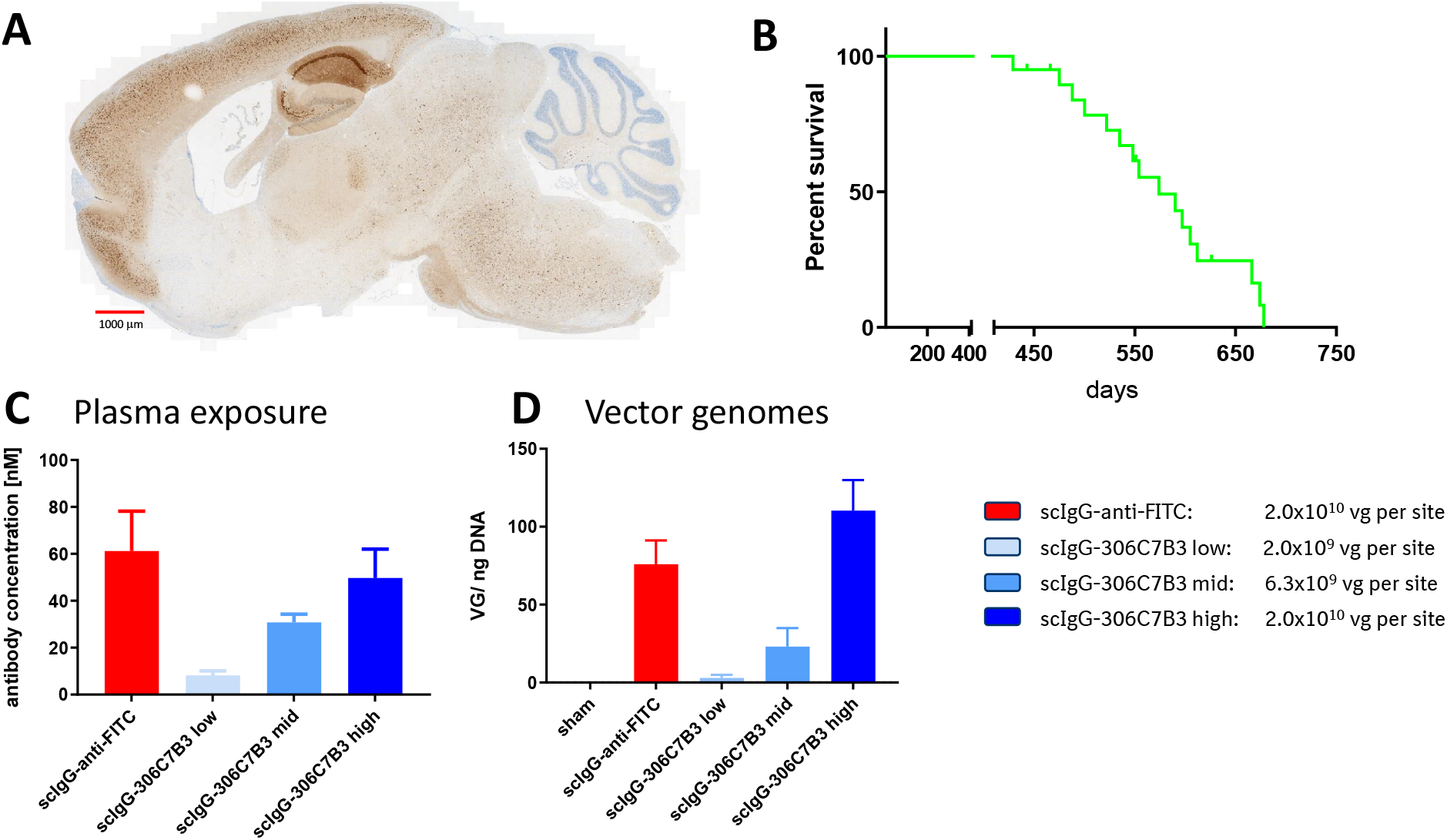
In vivo model to assess functional activity of the scIgG-306C7B3. ***A***, Sagital section of the brain of a (Thy-1)-[A30P]-hα-synuclein mouse overexpressing mutant A30P α-synclein under the control of the neuronal Thy-1 promoter euthanized due to a loss of righting reflex. Shown is a staining for pSer129-positive α-synuclein aggregates. The majority of aggregates are found in the hindbrain regions and areas of the Medulla oblongata. See supplemental figure 5 for more details. ***B***, Kaplan-Meier plot of non-treated (Thy-1)-[A30P]-hα-synuclein mice demonstrating age-dependent survival based on loss of righting reflex. ***C***, Animals received bilateral stereotactic inoculation of each 1 μl of AAV2HBKO preparations carrying transgenes coding for either scIgG-306C7B3 or scIgG-antiFITC control antibody at the indicated viral titers (left). Animals were inoculated at the age of 12 months and analysed for plasma exposure after loss of righting reflex (mean plus SEM) analysed with MSD-based ELISAs corresponding to the analyzed samples. No CSF samples had been taken from the actual in vivo study analysing the functional activity of scIgG-306C7B3 for practical reasons. ***D***, Post-mortem analysis of brain tissue for transduction efficacy (vg = vector genomes per ng of genomic DNA), demonstrating dose-dependent transduction (mean plus SEM). Sham = non-inoculated controls.

A pilot experiment (Figure 8) was performed to confirm CNS exposure to scIgG-306C7B3 as well as scIgG-anti-FITC (control, supplementary Figure 2B) using a dose of 2.0×10^10^ vector genomes of AAV2HBKO per site. Plasma as well as cerebrospinal fluid scIgG-306C7B3 and scIgG-anti-FITC content were analyzed at days 28, 42 and 56 post-inoculation by MSD-based ELISA, and confirmed long-term, chronic exposure. Plasma exposure presumably derived from continuous washout of CSF due to lymphatic drainage was in the range of 50 – 100 nM (Figure 8A and 8B). Even more importantly, strong CSF scIgG-306C7B3 and scIgG-anti-FITC content in the single digit nanomolar range was observed. The ratio between plasma and central CSF scIgG-306C7B3 and scIgG-anti-FITC levels was calculated to be approximately 12.6 to 1, a value strongly deviating from the ratio achievable after peripheral administration of antibodies in patients alone (approx. 330 to 1, Jankovic et al., 2018) or from the naturally observed ratio for antibodies in the healthy situation (Reiber, 2001).

**Figure 8.**
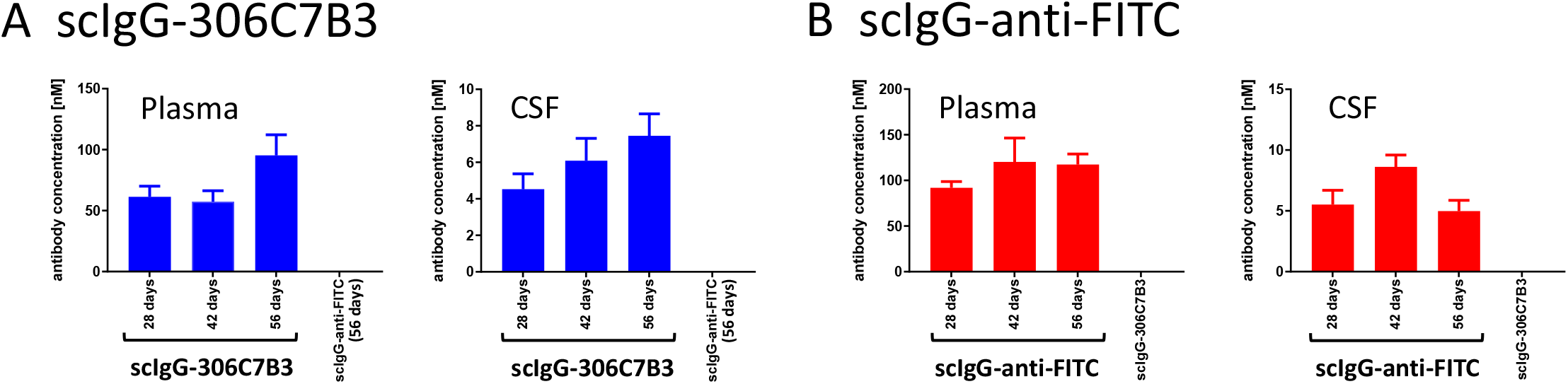
Pilot pharmacokinetic study to determine scIgG-306C7B3 and scIgG-anti-FITC exposure. (Thy-1)-[A30P] hα-synuclein mice were unilaterally inoculated with 2.0×10^10^ vector genomes into the striatum with either AAV2HBKO-scIgG-anti-FITC or AAV2HBKO-scIgG-306C7B3. Plasma as well as CSF fluid were analysed in transduced animals after 28, 42 and 56 days (separate cohorts for each individual time point). ***A***, Plasma exposure measuring 306C7B3 with a MSD based ELISA. ***B***, Central exposure in cerebrospinal fluid (CSF) taken at the indicated timepoints. Analysis as in A. Taking all data together, an approximate ratio of plasma to CSF exposure of 12.6 : 1 has been calculated for scIgG-306C7B3. Data presented as mean plus SEM (n=8 animals per group).

Based on this pilot experiment, three doses for AAV2HBKO scIgG-306C7B3 (2.0×10^9^ vg, 6.3×10^9^ vg and 2.0×10^10^ vg per site, bilateral striatal inoculation) were selected for the functional in vivo characterization of scIgG-306C7B3. The control construct – scIgG-anti-FITC – was bilaterally inoculated at the high dose of 2.0×10^10^ vg only. In our hands, this animal model never presents with central pSer129-positive α-synuclein pathology if analyzed below 14 months of age (Scudamore and Ciossek, 2018; and data not shown). Thus, animals at an age of 12 months were selected for initiating treatment via AAV transduction. Only animals with clear symptoms corresponding to α-synucleinopathy within the CNS (loss of righting reflex) were included in the following analysis (see supplementary table 1 for details on all animals initially treated and reasons to exclude individual animals).

Figure 7C shows the dose-dependent antibody exposure achieved in this experiment based on the analysis of the final bleed, corresponding to extrapolated CSF exposures of 0.7 nM, 2.5 nM and 3.9 nM for the low, mid and high doses, respectively (based on CSF exposure data from the pilot experiment, Figure 8). Efficient, dose-dependent transduction was confirmed for all animals based on vector genomes present in postmortem brain tissue (Figure 7D).

The functional effect of scIgG-306C7B3 treatment in this model of AAV-mediated antibody expression based on increased survival is shown in Figure 9. Comparing the non-treated cohort with the scIgG-anti-FITC transduced cohort, no change in survival time could be observed between untreated (Thy-1)-[A30P]-hα-synuclein mice and scIgG-anti-FITC transduced animals (p=0.47, Figure 9A). This argues for the absence of overt detrimental effects of AAV2HBKO mediated antibody production in the murine striatum. Further comparisons were made towards scIgG-anti-FITC treated animals only to account for potential minor detrimental effects due to viral transduction as well as the consequences of the surgical procedure of stereotactic inoculation into aged mice. Comparison of low and mid doses of AAV2-HBKO coding for scIgG-306C7B3 to the high dose of control AAV2HBKO scIgG-anti-FITC (Figure 9B) revealed no statistically significant changes in overall survival of treated animals. Interestingly, initial survival seems to be prolonged for about 50 days, although none of the scIgG-306C7B3 treated animals survived longer than the control scIgG-anti-FITC treated animals.

**Figure 9.**
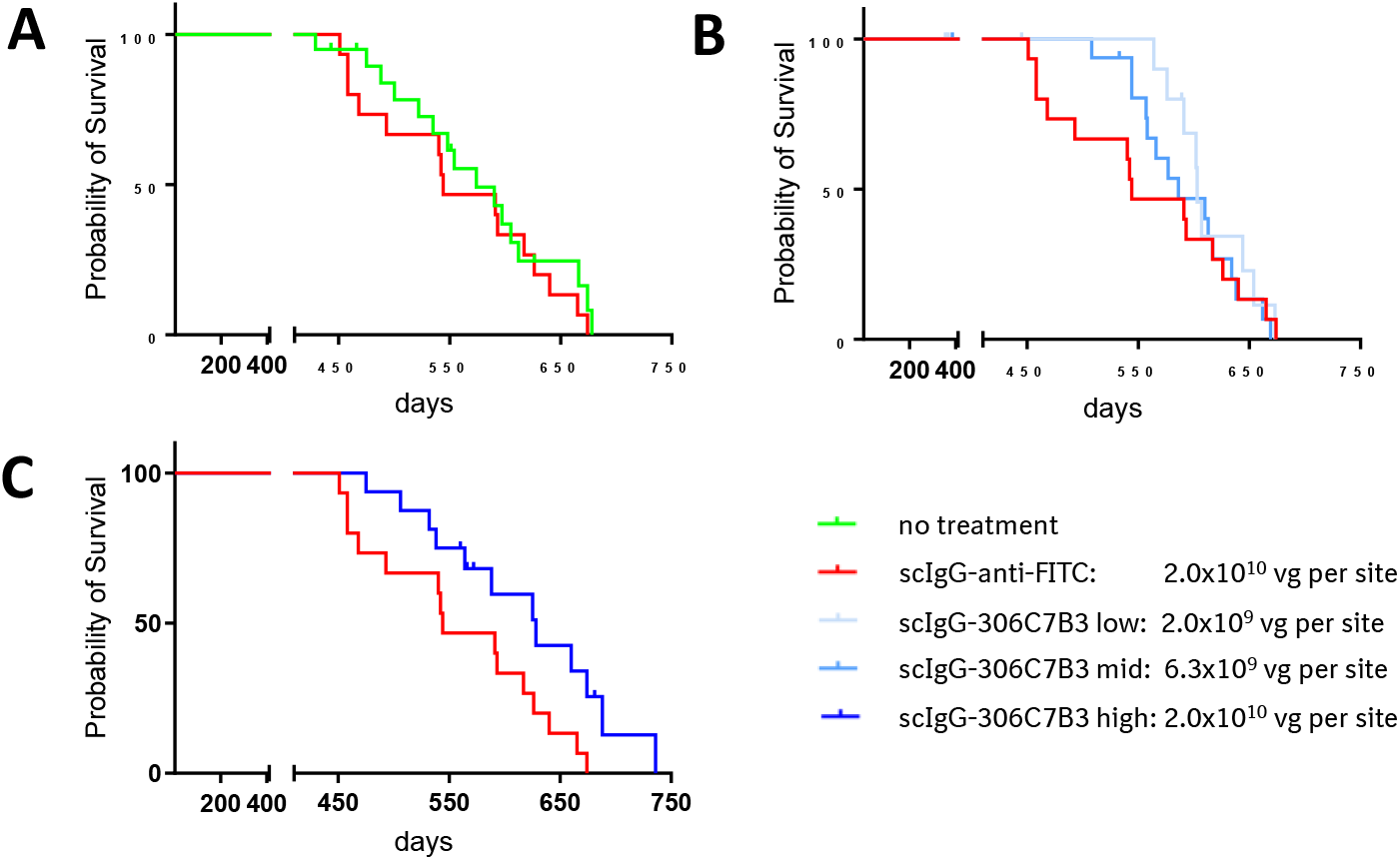
In vivo functional activity of scIgG-306C7B3 based on prolonged survival of AAV2-HBKO treated (Thy-1)-[A30P]-hα-synuclein mice. ***A***, Comparison of non-treated with control antibody treated (scIgG-anti-FITC) animals. No detrimental effect of the AAV-treatment could be observed (p=0.47). ***B***, Low and mid doses of AAV2HBKO coding for scIgG-306C7B3 demonstrate delayed initial mortality but no overall increase in survival (p=0.99 and 0.71, respectively). ***C***, Significant increased survival observed in high dose AAV2HBKO-scIgG-306C7B3 compared to control antibody treated animals (p=0.03; comparison towards non-treated animals p=0.09).

The survival of animals exposed to the high AAV2HBKO scIgG-306C7B3 dose was markedly greater than that of control scIgG-anti-FITC transduced animals (Figure 9C, p=0.03). Comparison to control animals (not inoculated with viral preparations) also showed a clear increased survival, but just missed statistical significance (p=0.09). Plasma scIgG-306C7B3 levels reached 49.7 nM at the high dose of AAV2-HBKO scIgG-306C7B3 (Figure 7C). These observations indicate that exposure of the CNS to a neutralizing, anti-aggregate specific α-synuclein antibody could block disease progression in this established animal model for α-synucleinopathies.

## Discussion

As pointed out in the introduction, one widely accepted model accounting for the temporal and spatial progression of pathology in α-synucleinopathies relies on prion-like spread of α-synuclein aggregates with seeding propensity between neurons. 306C7B3 has been initially developed based on the idea of blocking such prion-like spread of α-synuclein seeds in MSA, DLB or PD patients by passive immunization.

Successful therapeutic application of antibodies or any other larger biological entity in synucleinopathies or other neurodegenerative diseases where prion-like spreading occurs must fulfill several important requirements. α-synuclein aggregates with seeding propensity must be spatially accessible to antibodies that prevent their binding to and uptake into non-diseased neuronal or glial cells. The binding moieties of such antibodies should bind efficiently to epitopes present on α-synuclein seeds. It should not bind with high affinity to the monomeric form of the protein (e.g. as observed for PRX002 in a clinical trial (Schenk et al., 2017). The antibodies should further be devoid of off-target binding. A critical point that must be addressed as well is the achievable exposure level in patients. The central nervous system is known to represent an immune-privileged organ shielded from the peripheral environment by the blood-brain barrier that severely limits antibody access to the human brain. This has for example been recently confirmed in the clinical trials for PRX002 (Jankovic et al., 2018) and BIIB054 (Brys et al., 2019) for PD patients.

We describe here the characteristics of a novel anti-α-synuclein antibody exhibiting high affinity toward aggregated forms of this protein and poor, if any, binding to monomeric α-synuclein. In surface plasmon binding experiments the antibody displayed a very low off-rate with an overall calculated affinity of around 15 pM towards pre-formed fibrillar α-synuclein. Despite exhibiting recognition towards a relatively large amino acid stretch within α-synuclein C-terminal end, the PEPperPrint epitope analysis suggests that the epitope most recognized by 306C7B3 spans residues 133-139 (Figure 2). We therefore conclude that 306C7B3 recognizes a specific conformation that the α-synuclein C-terminal end adopts within its fibrillar form. We further conclude that the conformation 306C7B3 recognizes is not affected by Ser 129 phosphorylation (Figure 1).

306C7B3 binds with high affinity to structurally distinct fibrillar polymorphs and on fibrillar assembly α-synuclein oligomeric species. We conclude from this observation that the α-synuclein C-terminal end adopts the same conformation within all fibrillar species (Figure 3). More importantly, strong staining of human pathological α-synuclein aggregates in patient samples suffering from MSA was demonstrated. No binding to pathological protein deposits in other neurodegenerative diseases like Alzheimer’s Disease, Frontotemporal Dementia or Progressive Supranuclear Palsy, all known not to carry α-synuclein deposits, was observed. This suggests high specificity of 306C7B3 for pathological α-synuclein aggregates (Figure 4 and 5).

306C7B3 had been generated by classical immunization with *in vitro* aggregated, purified human α-synuclein, not carrying post-translational modifications (PTMs). Human α-synuclein is phosphorylated, ubiquitinated, acetylated and nitrated but the only residues that are subject to phosphorylation or nitration are S129, Y133 and Y136 within the α-synuclein C-terminal end (reviewed by Schmid et al., 2013), with pSer-129 modification being present in about 90% of the α-synuclein derived from pathological deposits (Anderson et al., 2006). Human α-synuclein is also subject to truncation at residues 133 and 134. As discussed in a recent review by Sorrentino and Giasson (2020), due to several technical limitations the fraction of truncated α-synuclein in postmortem samples can only be roughly estimated and seems to be in the range of ∼15-30% of total deposited α-synuclein with the majority of α-synuclein detected via MS-based analysis corresponding to the full-length protein, with abundancies of 0.23 to 0.26 of 1–122, 1–135, and 1–119 truncated forms compared with full-length 1–140 α-synuclein (set to 1.00; Kellie et al., 2014). Our immunohistochemical analysis of patient material argue for strong binding of 306C7B3 to pathogenic α-synuclein deposits despite PTMs and partial truncations.

It remains unclear which form of α-synuclein drives pathology, especially regarding the assumed pathological seeds underlying the spatial and temporal disease progression in patients. The fact that 306C7B3 strongly stains pathological deposits in postmortem brain samples argues for its potential to binds such seeds, pointing to the possibility that only a minor fraction of the α-synuclein in these seeds is C-terminally truncated.

As indicated previously, sufficient central exposure with therapeutic antibodies is key for treatment efficacy and this is severely limited due to the blood-brain barrier preventing significant transfer of peripheral antibodies into the brain (Reiber, 2001). This has been shown in preclinical animal models of neurodegeneration like e.g. Alzheimer’s Disease (Bien-Ly et al., 2015), but also seen in recent clinical trials of passive immunization against α-synuclein (Jankovic et al., 2018: Brys et al., 2019). We have previously performed a passive immunization study with a similar antibody. Here, up to 30 mg/kg of the therapeutic antibody was applied twice per week via intraperitoneal application for 15 weeks into (Thy-1)-[A30P]-hα-Synuclein mice. Pathology was induced by prior stereotactic bilateral inoculation of hα-synuclein PFF into the striatum, based on the observed pathological spread of disease described by Luk et al. (2012b). Although high plasma exposure in the single digit-micromolar range was achieved, no significant treatment efficacy could be observed (data not shown). A similar study approach in (Thy-1)-[A30P]-hα-Synuclein mice (Lindström et al., 2012), albeit not applying PFF-inoculation mediated pathology induction, also reported no significant effects on fibrillar forms of α-synuclein in insoluble fractions of the brain or spinal cord. Interestingly, the authors reported a trend towards increased survival of treated animals, which they attributed to a reduction in soluble and membrane-associated protofibrils in the spinal cord but not in the brain (Lindström et al., 2012).

Failure to achieve efficacy has been observed also in clinical studies on passive immunization in e.g. Alzheimer’s Disease trials (Nimmo et al., 2021), as well as recently in a PD trial using an anti-α-synuclein antibody (BIIB054/Biogen, trial discontinuation due to failure to achieve primary and secondary endpoints in Phase II clinical testing in PD patients; press release Biogen 2021). We therefore decided to explore AAV-based antibody expression, a gene therapy-based approach driving antibody expression directly within the tissue of interest (Fuchs et al., 2016). These approaches have been tested so far mostly in diseases requiring peripheral exposure (Robert et al., 2016; Balazs et al., 2012), but have been reported also regarding central transduction mediating antibody exposure within the brain (Liu et al., 2016; Ising et al., 2017; Fukuchi et al., 2006). At least one trial targeting HIV transmission by intramuscular application also reached clinical stage (Phase I clinical testing; Priddy et al., 2021), albeit facing problems regarding sufficient transgene expression.

The expression of secreted antibodies within the CNS will not only expose successfully transduced neurons to the therapeutic principle but also allow these antibodies to be present within the ISF as well as the CSF. The glympathic pathway, a convective influx from CSF into paravascular spaces mostly driven by aterial pulsatility (Needergard, 2013), will distribute these antibodies throughout the whole brain including the spinal cord (Iliff et al., 2012 and 2013), ensuring widespread central distribution even into non-transduced brain areas. CSF is constantly cleared via lymphatic pathways into the peripheral circulation (Sakka et al., 2011; Louveau et al., 2015), explaining the observed plasma exposures in our experiment.

Transduction with AAV2HBKO drove dose-dependent plasma exposure with scIgG-306C7B3 of up to 49.7 nM with a calculated CSF exposure of 3.9 nM, well in line with the experimental approach of central antibody expression combined with clearance into and accumulation in the peripheral blood circulation. Survival was analyzed to circumvent the variability in onset and aggregate load in the (Thy-1)-[A30P]-hα-Synuclein model (Freichel et al., 2007), with the loss of the righting reflex of affected animals as criterion closely correlated to disease progression (Freichel et al, 2007; Scudamore and Ciossek, 2018). No detrimental effects of neuronal expression of our control scIgG-anti-FITC antibody could be observed, demonstrating the general feasibility of such an approach. Interestingly, the low and mid dose of AAV2HBKO scIgG-306C7B3 showed signs for efficacy without overall increased survival, with efficacy indicated at younger age (Figure 9B). Statistically significant increased survival was only observed with the high AAV2HBKO dose, correlating with the higher overall plasma and CSF exposure. This data indicates that an initial protective effect of 306C7B3 might be overwhelmed by disease progression at later timepoints. It is important to consider that the (Thy-1)-[A30P]-hα-Synuclein model strongly overexpresses mutant [A30P]-hα-synuclein, driving disease induction and progression without additional manipulations like PFF inoculation. Available data suggests that prion-like seeds of α-synuclein are directed to the endolysosomal pathway for degradation and can induce pathological intracellular aggregation only upon endosomal escape (Karpowicz et al., 2017; Sacino et al., 2016; Flavin et al., 2017). Aging is a major risk factor for many neurodegenerative diseases including PD. Evidence exists from preclinical models that the lysosomal activity (measured as macro-autophagy and chaperone-mediated autophagy activity) decreases with aging, although data for the human brain is scarce (Loeffler, 2019). We hypothesize that our reported, dose-dependent effects on survival might reflect the ongoing decline of lysosomal activity in the murine brain enabling α-synuclein seed degradation, causing an age-dependent increase in prion-like seeds ultimately driving disease progression. Lower exposure with 306C7B3 might be initially sufficient to diminish prion-like disease spreading but this effect is eventually overwhelmed upon further aging. Although disease progression finally occurs also within the cohort of animals treated with the high dose of AAV2HBKO-scIgG-306C7B3, it appears that the higher CSF exposure with 306C7B3 is sufficient for increased survival. Additional experiments are required to further strengthen this hypothesis.

Several antibodies targeting α-synuclein have been tested in models for α-synucleinopathies to prevent PD-like pathology (see recent reviews by Folke et al., 2022; Vijayakumar and Jankovic, 2022; Castonguay et al., 2021), albeit with mixed outcomes. Many approaches went for passive immunization with antibody delivery via intraperitoneal application. Most studies utilized a preventive setting, in which the protective anti-α-synuclein antibody was delivered prior to active disease induction either upon PFF inoculation (Weihofen et al., 2019), potentially compromised by exposure of the inoculated PFFs to the inactivating antibodies during the surgical procedure disrupting the blood brain barrier. In a prophylactic treatment using the (Thy-1)-[A30P]-hα-Synuclein model (Nordström et al., 2021), significantly weaker effects were reported with pathology reduction mainly analyzed based on reduced pSer129 staining. The effect of the neutralizing antibody was not dose-dependent and potential interference with binding of the utilized pSer129 antibody was not analyzed (epitope of mAb47/ABBV-0805 corresponds to amino acids 121-127 of hα-synuclein, close to the pSer129 site; Nordström et al., 2021). MEDI1341, another α-synuclein antibody, has only been analyzed regarding reduction of staining for human α-synuclein without testing for effects on pSer129 α-synuclein staining, indicative of insoluble α-synuclein aggregates. Questions towards the MEDI1341-mediated masking of the utilized Syn-1 IHC staining remain (Schofield et al., 2019). Henderson et al. (2020) went for a therapeutic application of Syn9048 in wildtype mice with PFF inoculation-induced pathology, however no significant improvements in behavioral testing and only minor improvements in α-synuclein pathology in some specific brain areas were found, combined with an attenuation of dopamine reductions in the striatum. Recently, Choi et al. (2022) described efficacy of conformation-specific α-synuclein antibodies upon peripheral administration in the Line61 mThy1-α-syn Tg model. The underlying reason for this discrepancy to our previous, unpublished data with passive immunization in the (Thy-1)-[A30P]-hα-Synuclein model remains unclear, however the Line61 model displays significantly weaker overall pathology in the brain. The same is true for the reported protective effects of active immunization with UB-312 in the Thy1SNCA/15 mouse model, with improvements in behavior scores and reductions in the levels of α-synuclein oligomer levels 15 weeks after immunization. However, the main protective effects are observed in the colon in this model, potentially hinting towards a main effect on diminishing spreading of pathology from the colon to the CNS (Nimmo et al., 2022).

Quite similar approaches to the one described in this publication have been followed by Chen et al. (2021) and Butler et al. (2022) with AAV-mediated expression of fibril-specific α-synuclein binding nanobodies within the CNS. However, these approaches target intracellular α-synuclein by expression of single-chain intrabodies that are not secreted and that exert protective effects only within actually transduced cells. Such an approach is currently not translatable to clinical use requiring targeting the whole human brain being approximately 3000 times larger than the murine brain.

To the best of our knowledge, this is the first description of AAV viral vector mediated expression of a secreted, full-length anti-α-synuclein antibody in a mouse model for α-synucleinopathies resulting in prolonged survival. Based on its unique properties, we believe that this approach of central, AAV-mediated expression of 306C7B3 presents an interesting treatment option for PD or MSA. In addition, application of 306C7B3 for diagnostic purposes deserves further consideration.

## Author contributions

CK, TL, SL, RM, MN, DW, FI and TC designed the studies. TC wrote the manuscript. MD, DB, PR and JS performed the actual experiments. TS, MN and BS analyzed the immunohistopathology.

## Data Availability

Data supporting the findings of this study (high resolution IHC figures, raw data) are available from the corresponding author upon reasonable request.

---

**Supplementary Figure 1.**
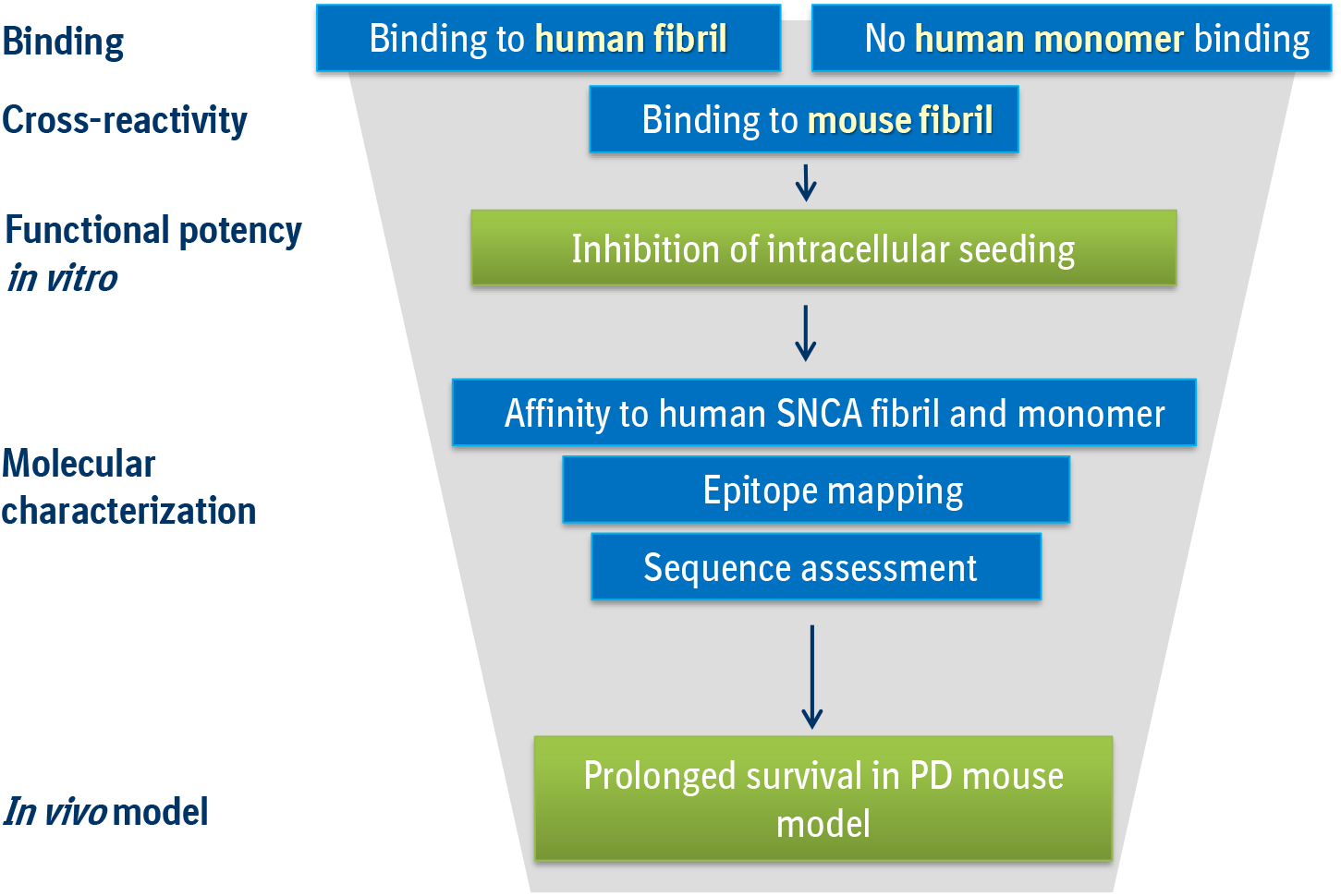
Screening cascade employed to identify fibril-specific anti-α-synuclein antibodies. Hybridoma supernatants were screened by ELISA for binding activity against fibrillar human and mouse α-synuclein. Supernatants of hybridomas negative for binding towards monomeric human α-synuclein were tested in a cellular assay assessing their capacity to block intracellular seeding (SH-SY5Y cells stably overexpressing mutant A53T human α-synuclein). Selected clones were sequenced and underwent an epitope mapping (PepperPrint assay). Finally, 306C7B3 was tested in the described in vivo Parkinson’s Disease (PD) model in (Thy-1)-[A30P]-hα-Synuclein mice.

**Supplementary Figure 2A.**
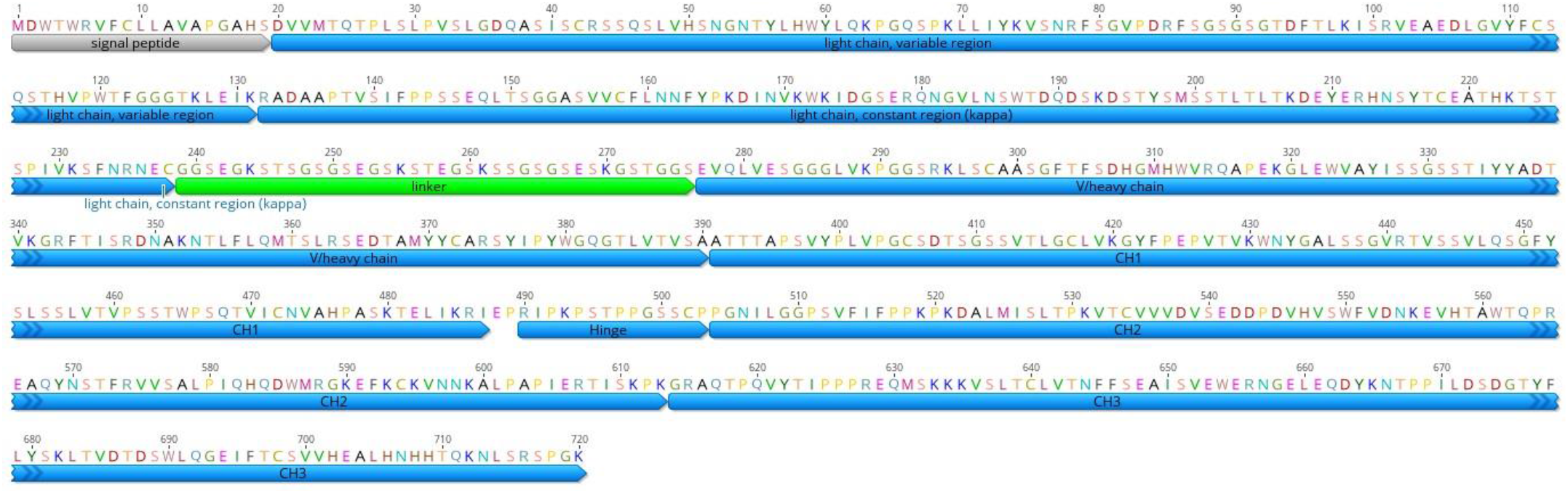
Amino acid sequence of murine scIgG-306C7B3. Light and heavy chains are encoded within one single open reading frame due to the use of a linker sequence (green). The human IgG1 signal peptide was added to the primary antibody sequence to allow for efficient secretion of the scIgG from transduced cells (signal peptide).

**Supplementary Figure 2B.**
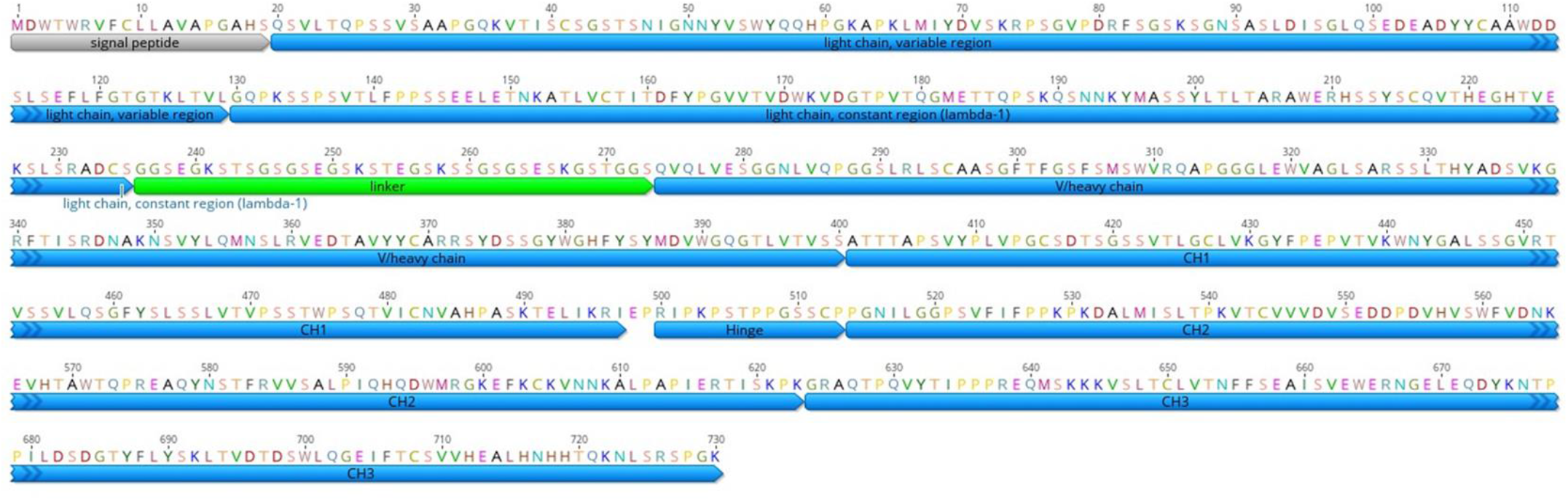
Amino acid sequence of murine scIgG-anti-FITC. Light and heavy chains are encoded within one single open reading frame due to the use of a linker sequence (green). The human IgG1 signal peptide was added to the primary antibody sequence to allow for efficient secretion of the scIgG from transduced cells (signal peptide). The anti-FITC IgG sequence is derived from the scFv anti-FITC nanobody as described in Vaughan et al (1996).

**Supplementary Figure 2C.**
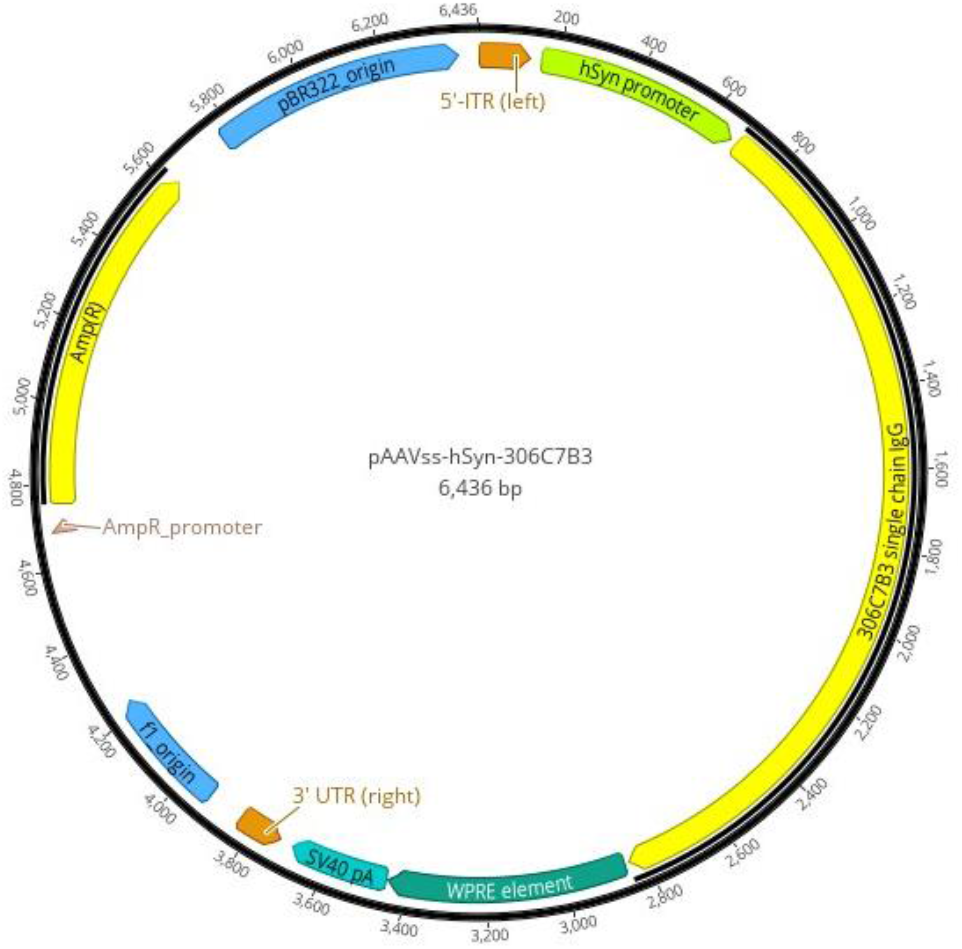
Plasmid map of pAAVss-hSyn-scIgG-306C7B3. The expression cassette – flanked by the two inverted terminal repeats of AAV2 required for efficient packaging into AAV capsids – contains the human synapsin promoter (hSyn promoter), the full scIgG-306C7B3 antibody sequence in a single open reading frame including an optimized Kozak sequence, a WPRE element and the SV40 polyadenylation signal (SV40 pA).

**Supplementary Figure 3.**
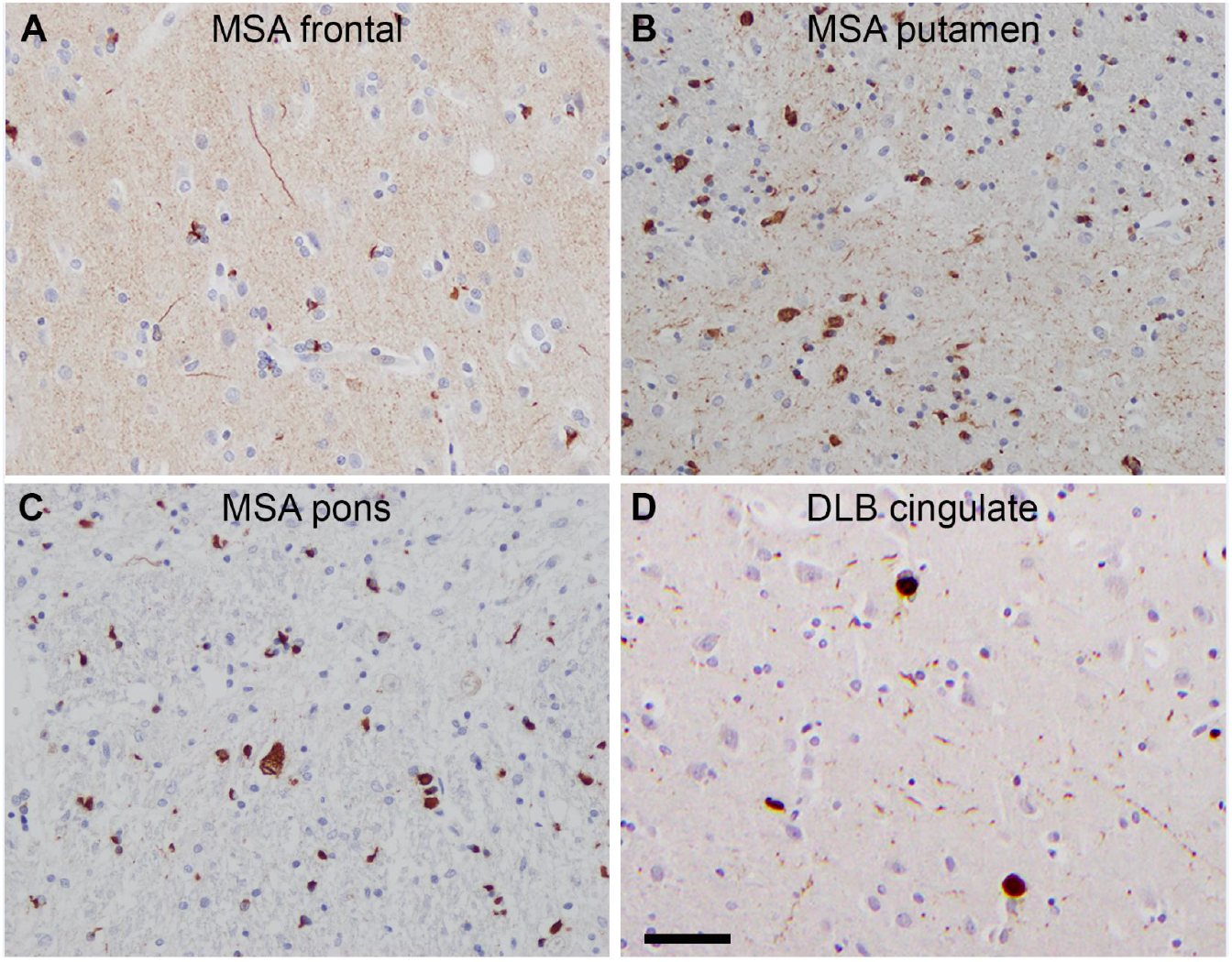
306C7B3 immunoreactivity in multiple system atrophy and Lewy body disease. Immunohistochemistry for 306C7B3 in formalin-fixed paraffin embedded section of human postmortem tissue of α-synucleinopathies. Robust immunoreactivity is observed for all types of inclusions in MSA including cortical and subcortical neuronal and oligodendroglial inclusions in frontal cortex (A), putamen (B), and pons (C). In Lewy body diseases, strong labelling of Lewy bodies and Lewy neurites is seen as illustrated in cingulate gyrus of dementia with Lewy body case (D). Scale bar 50μm (A-D).

**Supplementary Figure 4.**
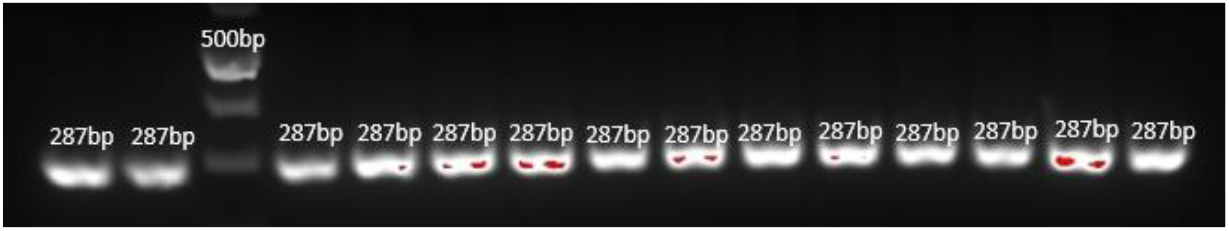
Confirmatory biochemical analysis of all animals involved in the in vivo study. Gel electrophoresis example showing the result of the genotype analysis via PCR. All animals in the study were shown to be homozygous for the A30P α-synuclein transgene as indicated by the amplified 287 bp band from genomic DNA isolated from each individual animal.

**Supplementary Figure 5.**
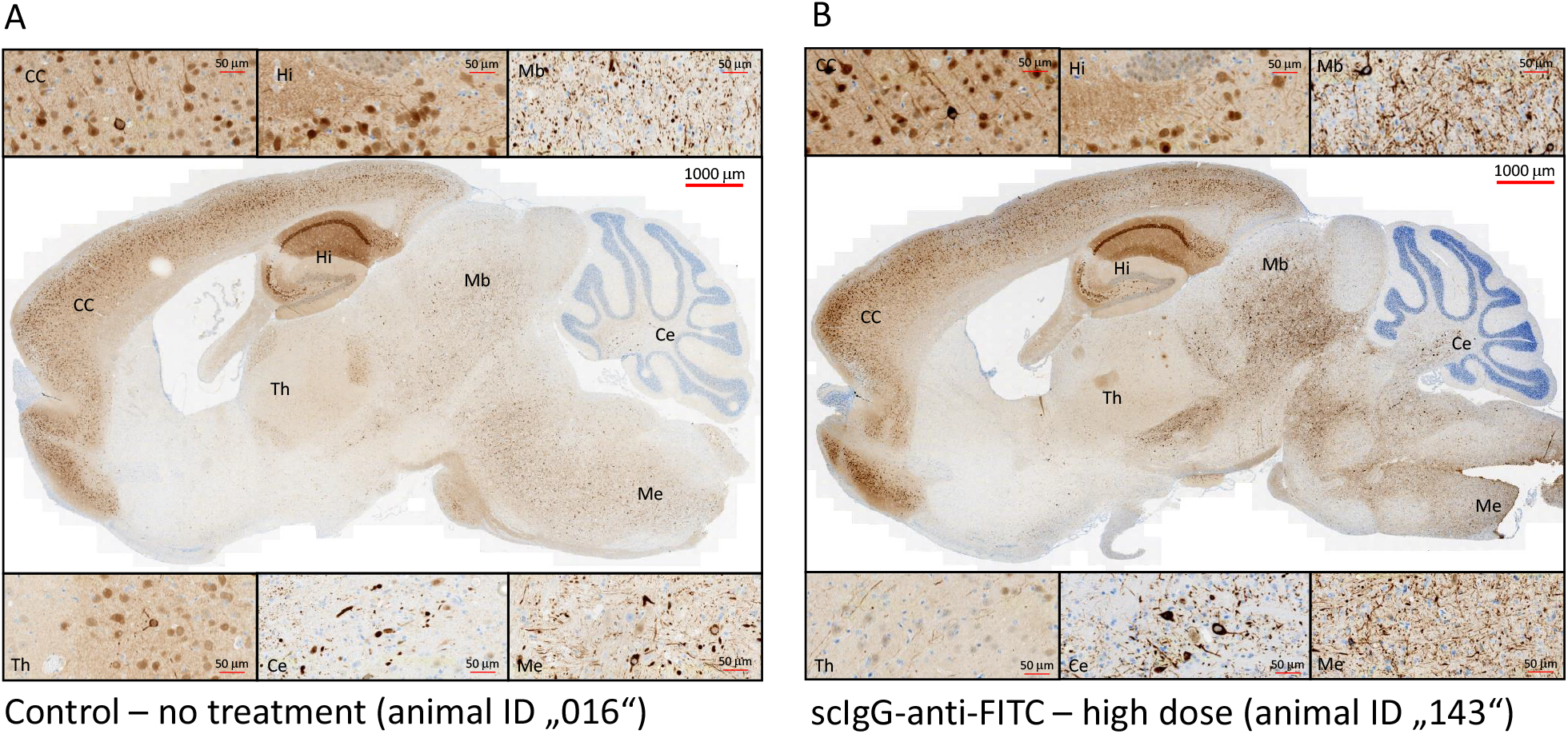
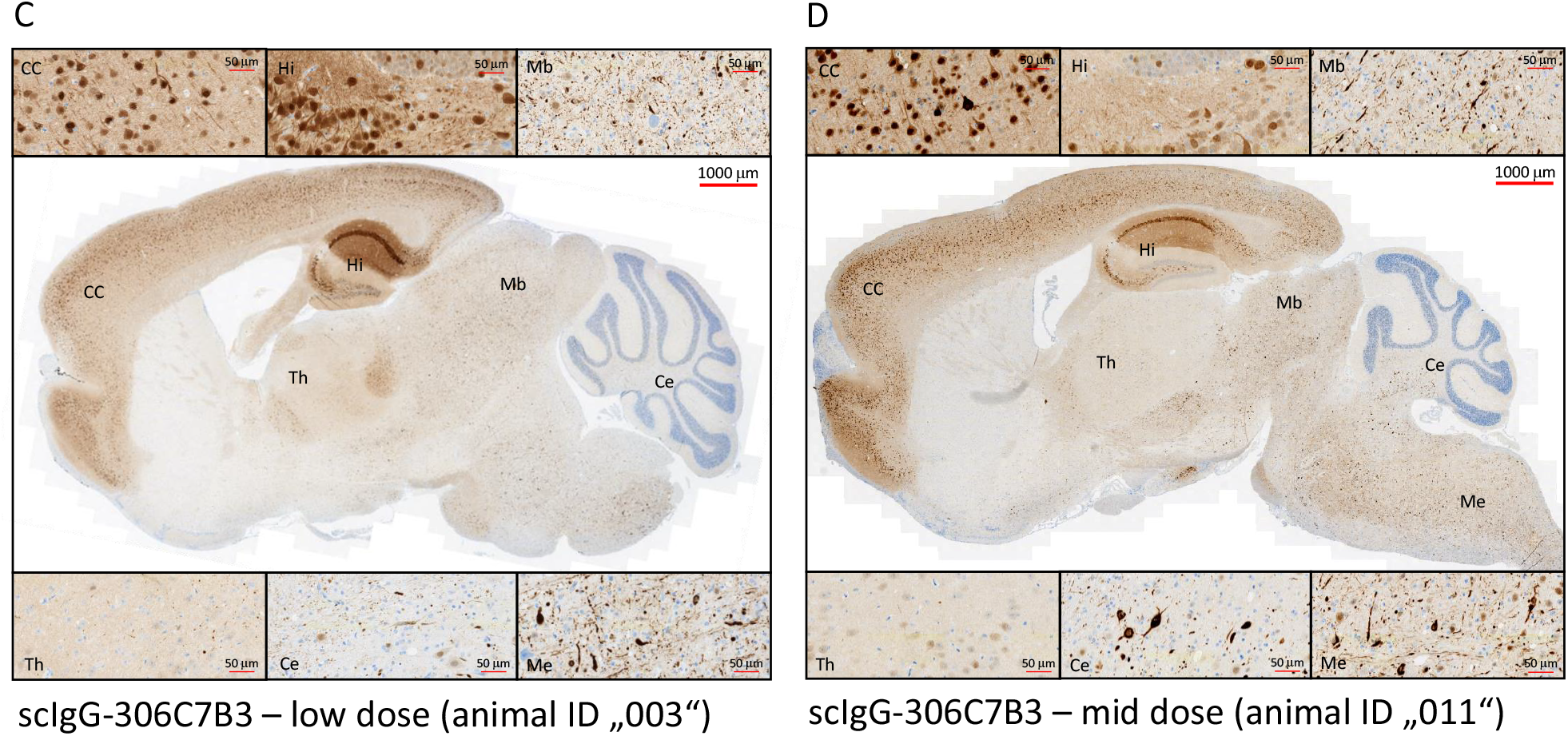
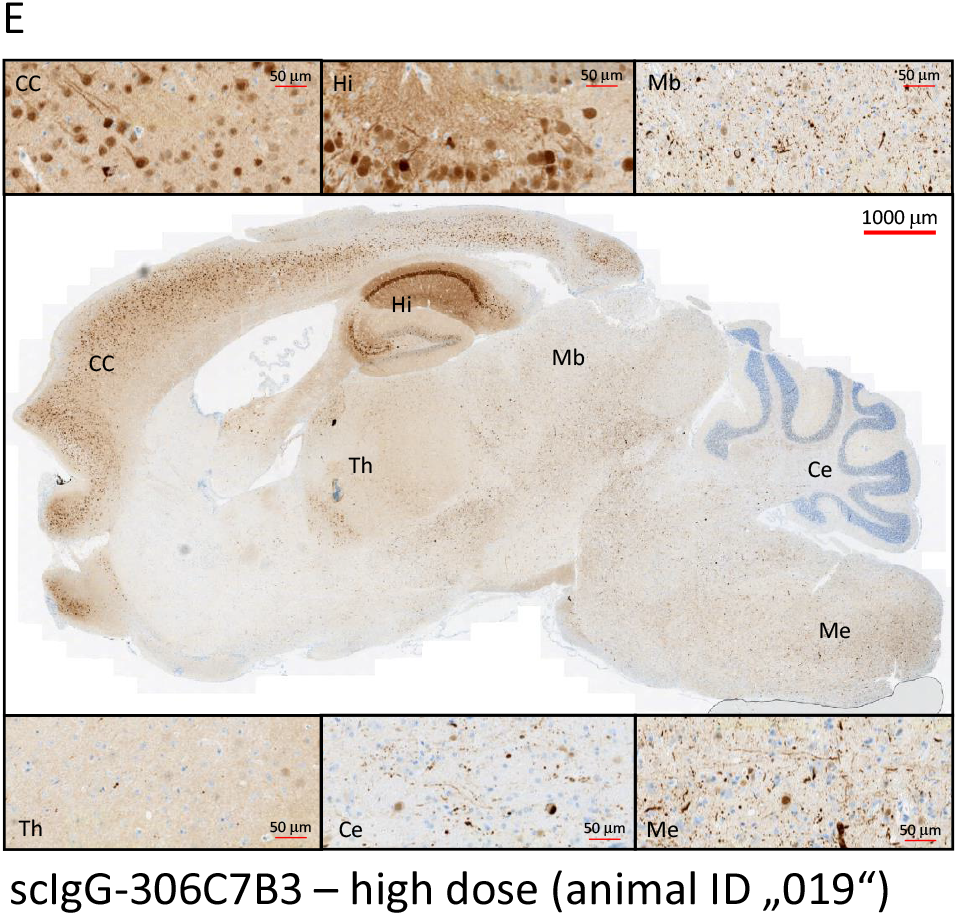
Confirmatory IHC analysis of the animals involved in the *in vivo* study. Formalin-fixed brain tissue from all animals included in the study were tested via pSer129-α-synuclein IHC for strong pathology, corresponding to the symptom of loss of righting reflex at time of euthanasia. Examples of the obtained stainings for representative animals from each cohort are shown. See supplementary table 1 for details on the individual animals. High magnification of the indicated areas are shown above and below the full brain images. CC: cerebral cortex, Hi: hippocampus, Mb: midbrain, Th: thalamus, Ce: cerebellum, Me: medulla.

