## Supplementary Table 1 for "AAV-mediated Expression of a Novel Conformational Anti-Aggregated α-Synuclein Antibody Prolongs Survival in a Genetic Model of α-Synucleinopathies"

| | | Gender | loss of righting reflex | age at sacrifice | |
|---|---|---|---|---|---|
| Group | animal ID | (m / f) | (yes / no) | (days) | comments |
| control | 8 | m | yes | 475 | |
| | 9 | f | yes | 605 | |
| | 10 | f | yes | 535 | |
| | 12 | f | yes | 574 | |
| | 13 | m | no | 466 | found dead in cage |
| | 14 | m | no | 551 | large wound in the back and at the ear, euthanized |
| | 15 | m | yes | 666 | |
| | 16 | f | yes | 522 | |
| | 17 | f | yes | 429 | |
| | 18 | f | yes | 597 | |
| | 19 | f | yes | 488 | |
| | 20 | f | yes | 590 | |
| | 111 | f | yes | 500 | |
| | 112 | f | yes | 548 | |
| | 172 | m | yes | 674 | |
| | 173 | m | yes | 612 | |
| | 174 | m | yes | 678 | |
| | 175 | f | no | 626 | rectum prolapse, euthanized |
| | 176 | f | yes | 554 | |
| | 177 | f | no | 575 | found dead in cage |
| | 178 | f | no | 443 | found dead in cage |
| high dose AAV2HBKO-scIgG-antiFITC 2.0E+10 vg | 1 | f | yes | 458 | |
| | 2 | f | yes | 468 | |
| | 3 | f | yes | 544 | |
| | 4 | f | yes | 593 | |
| | 5 | m | yes | 451 | |
| | 6 | m | yes | 617 | |
| | 7 | m | yes | 640 | |
| | 8 | m | yes | 493 | |
| | 143 | f | yes | 458 | |
| | 144 | f | yes | 665 | |
| | 146 | m | yes | 591 | |
| | 147 | m | yes | 540 | |
| | 168 | m | yes | 626 | |
| | 169 | m | yes | 674 | |
| | 170 | m | yes | 542 | |
| | 171 | m | no | - | euthanized shortly after surgery |
| low dose | 1 | f | yes | 576 | |
| | 2 | f | yes | 564 | |
| | 3 | f | yes | 591 | |

| | | | | | |
|---|---|---|---|---|---|
| AAV2HBKO-scIgG-306C7B3 2.0E+09 vg | 4 | f | no | - | general deterioration shortly after surgery |
| | 5 | m | no | 589 | self inflicted large scale scratches, euthanized |
| | 6 | m | yes | 607 | |
| | 7 | m | yes | 673 | |
| | 8 | m | no | - | general deterioration shortly after surgery |
| | 17 | m | yes | 603 | |
| | 18 | m | yes | 602 | |
| | 19 | m | no | - | general deterioration shortly after surgery |
| | 148 | f | no | 455 | vaginal prolapse, euthanized |
| | 149 | f | no | - | general deterioration shortly after surgery |
| | 150 | m | yes | 654 | |
| | 151 | m | yes | 644 | |
| mid dose AAV2HBKO-scIgG-306C7B3 6.3E+09 vg | 9 | f | yes | 566 | |
| | 10 | f | yes | 577 | |
| | 11 | f | yes | 613 | |
| | 12 | f | yes | 544 | |
| | 13 | m | yes | 508 | |
| | 14 | m | yes | 638 | |
| | 15 | m | no | 533 | rectum prolapse, euthanized |
| | 105 | m | yes | 669 | |
| | 152 | f | yes | 558 | |
| | 153 | f | yes | 634 | |
| | 154 | m | yes | 544 | |
| | 155 | m | yes | 586 | |
| | 164 | f | yes | 610 | |
| | 165 | f | yes | 557 | |
| | 166 | m | no | - | general deterioration shortly after surgery |
| | 167 | m | yes | 617 | |
| high dose AAV2HBKO-scIgG-306C7B3 2.0E+10 vg | 17 | f | yes | 538 | |
| | 18 | f | yes | 628 | |
| | 19 | f | yes | 564 | |
| | 20 | f | yes | 506 | |
| | 101 | m | yes | 660 | |
| | 102 | m | yes | 475 | |
| | 103 | m | yes | 736 | |
| | 104 | m | no | 572 | rectum prolapse, euthanized |
| | 139 | f | no | 566 | large, focal swelling at tail, euthanized |
| | 140 | f | no | 560 | large weeping wound at left leg, euthanized |
| | 141 | m | yes | 532 | |

| | | | | | |
|---|---|---|---|---|---|
| | 142 | m | yes | 588 | |
| | 159 | m | yes | 674 | |
| | 160 | m | no | - | general deterioration shortly after surgery |
| | 161 | m | no | 681 | general deterioration, no loss of righting reflex |
| | 162 | m | yes | 688 | |
| | 163 | m | yes | 625 | |

**Supplementary Table 1.**

Individual animal data for the in vivo functional study. Only animals with clear loss of righting reflex were included in the analysis. See comments for animals euthanized or found dead without loss of righting reflex.
